# Algal photosystem I dimer and high resolution model of PSI:plastocyanin complex

**DOI:** 10.1101/2021.08.30.458224

**Authors:** Andreas Naschberger, Laura Mosebach, Victor Tobiasson, Sebastian Kuhlgert, Martin Scholz, Annemarie Perez-Boerema, Thi Thu Hoai Ho, Andre Vidal-Meireles, Yuichiro Takahashi, Michael Hippler, Alexey Amunts

## Abstract

Photosystem I (PSI) enables photo-electron transfer and regulates photosynthesis in the bioenergetic membranes of cyanobacteria and chloroplasts. Being a multi-subunit complex, its macromolecular organization affects the dynamics of photosynthetic membranes. Here, we reveal a chloroplast PSI from the green alga *Chlamydomonas reinhardtii* that is organized as a homodimer, comprising 40 protein subunits with 118 transmembrane helices that provide scaffold for 568 pigments. Our cryo-EM structure identifies that the absence of PsaH and Lhca2 gives rise to a head-to-head relative orientation of the PSI-LHCI monomers in a way that is essentially different from the oligomer formation in cyanobacteria. The light-harvesting protein Lhca9 is the key element for mediating this dimerization. The interface between the monomers is lacking PsaH, and thus partially overlaps with the surface area that would bind one of the LHCII complexes in state transitions. We also define the most accurate available PSI-LHCI model at 2.3 Å resolution, including a flexibly bound electron donor plastocyanin, and assign correct identities and orientations of all the pigments, as well as 621 water molecules that affect energy transfer pathways.

## Main

A chloroplast PSI of green algae consists of the core complex and three antenna modules: inner belt, outer belt, and Lhca2:Lhca9 heterodimer, which together comprise 24 subunits ^1–4^. As a short-term light acclimation mechanism in response to fluctuating illumination and anoxia, the algal PSI additionally associates with two LHCII trimers ^5, 6^. Structural studies have shown that the oligomeric state of a chloroplast PSI is a monomer, due to the presence of the subunit PsaH, whereas in cyanobacteria structures of dimers ^7–10^ and trimers ^11^ were also reported. Cyanobacterial PSI oligomerizes via direct contacts between subunits PsaI and PsaL, however such an association has been ruled out for a chloroplast PSI due to structural constraints of PsaH presence that imposes an apparent rigidity ^12, 13^. Yet, recent structural studies of PSI from a chloroplast of a salt-tolerant alga suggested that its functional core may vary more than previously believed ^14^. Particularly, functional PsaH-free particles were found, thus showing a potential architectural plasticity of PSI in response to the ecological environment. On the macromolecular level, an atomic force microscopy analysis of a plant thylakoid membrane showed that when its architecture is altered upon transition from darkness to light, larger inter- membrane contacts are formed, leading to a reduced diffusion distance for the mobile electron carriers ^15^. The membrane architecture in dark- and light-adapted membranes contains ordered rows of closely packed PSI dimers, which are more abundant in the dark state ^15^. Similarly, closely associated PSI-LHCI complexes were detected in plants by negative stain electron microscopy ^16^, and dimers were found in a subpopulation of PSI from a temperature-sensitive PSII mutant alga ^17^. This suggests that reversible PSI dimer formation may have a physiological role in thylakoid membrane structure maintenance in chloroplasts. However, very little is known about PSI-LHCI dimers and information on their structures is lacking. In the absence of high resolution data, no evidence is available on composition, elements regulating and mediating dimerization, and how the arrangement would differ from the cyanobacterial counterparts.

### Structure determination

We grew *C. reinhardtii* cells containing a His-tag at the N-terminus of PsaB in low light and under anoxic conditions (see Methods). The thylakoid membranes were solubilised with *n-* dodecyl*-α-D-*maltoside (α-DDM), followed by affinity purification, crosslinking via the chemically activated electron donor plastocyanin (Pc) and sucrose density gradient centrifugation (Extended Data Fig. 1). Two PSI fractions were detected on the sucrose gradient, and 2D polyacrylamide gel electrophoresis (native/reducing 2D-PAGE) of isolated thylakoids indicated the presence of PSI dimers (Extended Data Fig. 2 and Supplementary Table 1). The heavier green band on the gradient was subjected to single-particle cryo-EM analysis (Supplementary Table 2). We used 2D classification to separate PSI dimers from monomers in a reference free manner, followed by 3D classification leading to a subset of 14,173 particles, which were refined to an overall resolution of 2.97 Å by applying C2 symmetry (Extended Data Fig. 3). PSI dimers were also found in 2D class averages in a dataset recorded from a sample without the use of crosslinker. Upon symmetry expansion, the resolution was further improved to 2.74 Å (Extended Data Fig. 3). The remaining 74,209 particles containing the monomer were refined to 2.31 Å resolution, representing a considerable improvement on the previously reported maps ^1–3^. A density corresponding to the bound electron donor Pc was found at the luminal side of both PSI forms, dimer and monomer.

### Overall structure

To derive a structure of the chloroplast PSI dimer, we first built an accurate model of one monomer using the 2.74 Å resolution map, and then fitted it into the cryo-EM density of the C2 refined dimer. Compared to the monomer, all but two core subunits (PsaH, PsaO) and one light-harvesting protein (Lhca2) are found in the dimer (Fig. 1, Supplementary movie 1). The structure contains 40 protein subunits, 398 chlorophylls *a*, 60 chlorophylls *b*, 56 beta-carotenes, 54 luteins, 2 violaxanthins, 2 neoxanthins, 4 phylloquinones, 6 iron-sulphur clusters, and 32 lipids (Fig. 1). In addition, two unaccounted densities corresponding to two loops in the stromal side of Lhca9 and PsaG could be interpreted in the dimer, due to the stabilization by the adjacent monomer (Extended Data Fig. 4). The first better-defined density is the Lhca9 loop region 132-153 which is stabilised due to a direct interaction with PsaL of the second monomer within the dimer. As a result, the area is closely packed with PsaG, and therefore also the PsaG loop region 63-77 is better resolved in the dimer (Extended Data Fig. 4). Finally, cofactors of Lhca9 could be modelled at the interface between the monomers. No density for the His-tag on PsaB could be detected. Overall, taking into account the challenges of modelling photosynthetic complexes ^18^, we were able to further improve the quality of the structure, resulting in the significantly better validation statistics, compared to the most recent cryo-EM ^17^ and X-ray crystallography studies ^19, 20^ (Supplementary Table 1). Thus, the current study represents the most complete reference model of PSI.

**Fig. 1:**
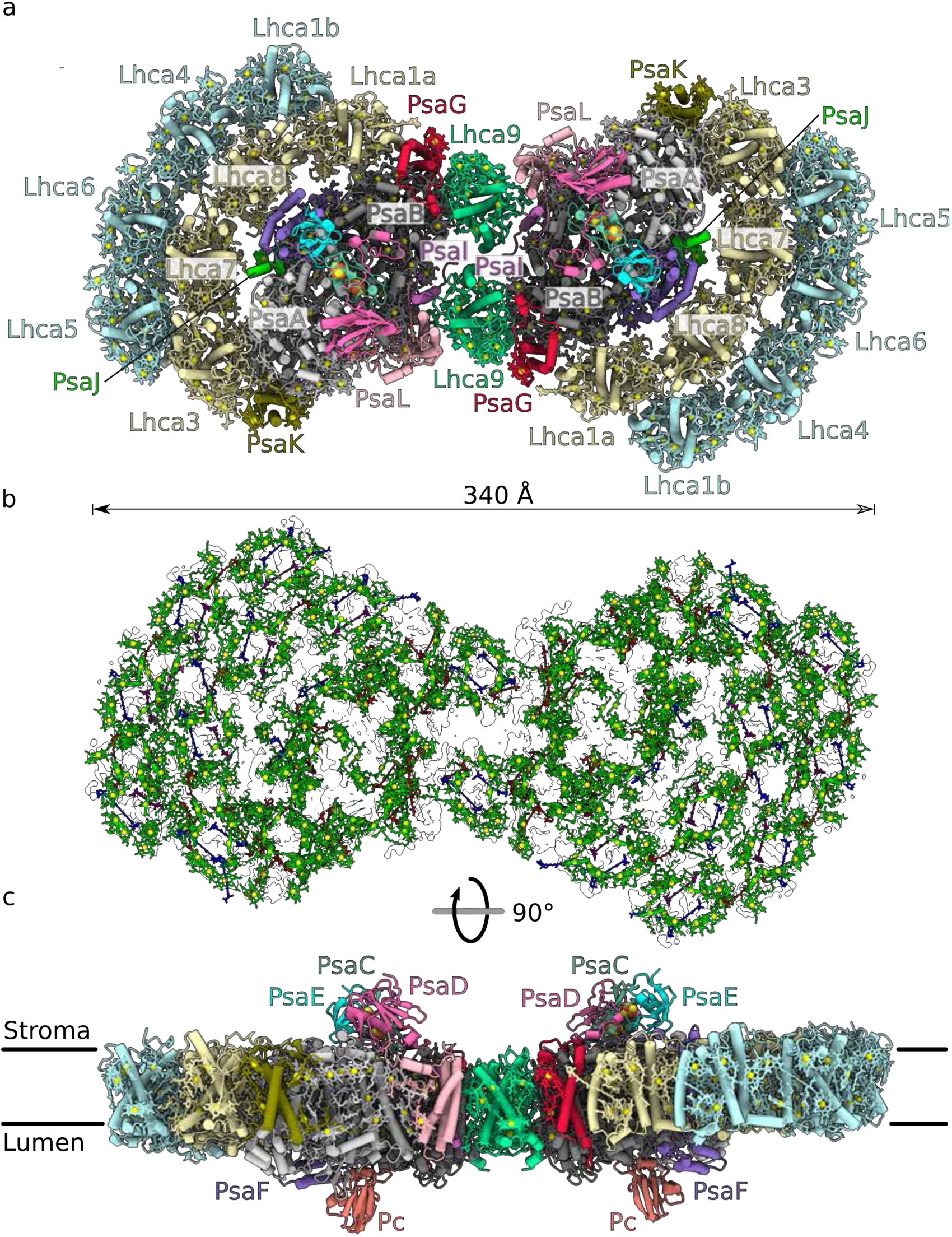
Overall structure of the PSI dimer. **a**, View of individual proteins from stroma. **b**, Arrangement of the pigments in the outline of the map: chlorophylls green (Mg yellow), luteins blue, beta-carotenes red, violaxanthin purple, and neoxanthin pink **c**, Overall view along the membrane.

### Structural basis for PSI dimerization

The structural basis for the algal chloroplast PSI dimerization is fundamentally different from cyanobacteria (Fig. 2a). In cyanobacteria, PSI dimerises via the stromal region of PsaL ^7–10^ and trimerises via the lumenal C-terminus of PsaL, assisted by PsaI ^11^. In our structure of the chloroplast PSI-LHCI dimer, neither PsaL nor PsaI interacts with each other between the neighbouring units. Instead, PsaH that normally preserves a monomer is not present, and Lhca9 with its associated cofactors acts as a symmetrical linker between the monomers, highlighting the importance of the light-harvesting antenna proteins for regulation of the macro-organisation. Lhca9 is distinct among the light-harvesting proteins in our structure due to a truncated loop between helices A and C, and lack of the associated chlorophyll ^6^. As a result, it contains the fewest chlorophylls among Lhcas (Supplementary Table 3). Based on this difference, we rationalised how Lhca9 allows for dimerization, as a longer AC-loop would clash with the neighbouring PsaB (Extended Data Fig. 5).

**Fig. 2:**
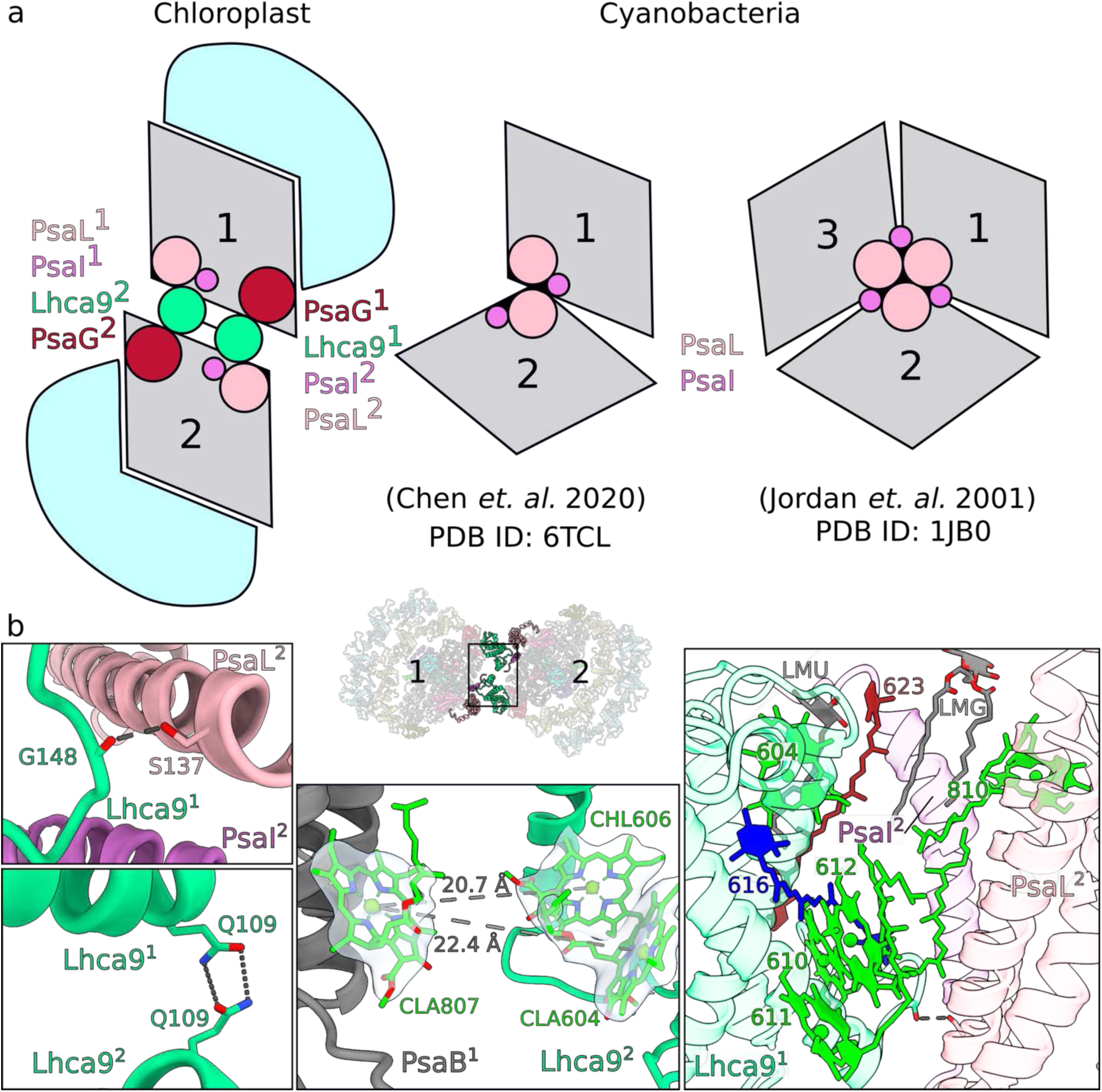
Dimerization of PSI. **a**, Schematic representation: chloroplast PSI-LHCI dimer associated via Lhca9, cyanobacterial PSI dimer (PDB ID: 6TCL) and trimer (PDB ID: 1JB0) associated via PsaI/L. The PSI core is grey, LHCI light-blue. **b**, The dimer interface is formed by: hydrogen bonds between PsaL and Lhca9, and between two Lhca9 copies (left); potential energy transfer paths between the two monomers (centre); pigments and lipids (right).

The two Lhca9 copies tether the PSI monomers in a head-to-head fashion, resulting in a 340-Å long structure (Fig. 1, Fig. 2b, Supplementary movie 1). They form interactions of four types covering the entire membrane span: 1) a hydrogen bond of the backbone carbonyl of G148 with S137 of PsaL in the stroma; 2) a hydrogen bond between the two Q109 of the Lhca9 copies; 3) hydrophobic contacts via coordinated cofactors in the membrane that include a newly modelled, beta-carotene 62 3 (N2 in nomenclature according to ref. ^6^), and five chlorophylls (604, 610, 611, 612, 810); 4) lipid-mediated hydrophobic interactions via monogalactosyl diglyceride LMG852 and LMU624. One acyl chain of lipid 852 associates with chlorophyll 810 from monomer-2, while the other acyl chain associates with beta-carotene BCR623 from monomer-1 (Fig. 2b). Thus, lipids contribute to the oligomerization of PSI, meaning that the membrane itself plays a role in the association. The finding that specific carotenes and lipids enable inter-molecular contacts that bridge the PSI monomers is of a particular interest, as it can only be detected by high-resolution structural studies. Similarly, a recent structure of the RC-LH1 dimer from *Rhodobacter sphaeroides* revealed a bound sulfoquinovosyldiacylglycerol that brings together each monomer forming an S-shaped array ^21^. The involvement of lipids in the oligomerization is consistent with the formation of supercomplexes in other bioenergetic membranes ^22–25^.

To solidify the structural observations, we engineered a *lhca9* insertional mutant having the His-tag at the N-terminus of PsaB and repeated the purification procedure in the same way as for the wild type. This time, no PSI dimer band could be found in the sucrose density gradient (Extended Data Fig. 6). Notably, Lhca9 is present in *Δlhca2*, while Lhca2 is absent from *Δlhca9* ^26^. Moreover, Lhca9 stably associates with PSI-LHCI after sucrose density gradient centrifugation of solubilized thylakoids isolated from *lhca2* insertional mutant (Extended Data Fig. 7).

### Implications of PSI dimerization

The specific interactions between the monomers are enabled due to unoccupied positions of PsaH and Lhca2. Since PsaH is also required for the lateral binding of LHCII to the PSI core in state transitions ^5, 6^, we next compared the structure of PSI-LHCI dimer to the state transition complex (Fig. 3). The superposition shows that Lhca9 from the neighbouring monomer is positioned in the membrane, where Lhca2 resides in PSI-LHCI-LHCII, and their three transmembrane helices would overlap with each other (Fig. 3b). The presence of the PsaH transmembrane helix is not compatible with the Lhca9^2^-associated cofactors CLA9, LMG852, BCR9 that extend from the neighbouring monomer in the dimer. In addition, the superposition shows that there would be a clash between PsaG and Lhca1 of the inner belt with one of the LHCII trimers, but not the other (Fig. 3b). Since Lhca2 and PsaH are absent, the structure of the algal PSI dimer would not facilitate LHCII binding at this position. However, our 2D-PAGE indicated a comigration of LHCII polypeptides with the dimer fraction, and therefore a structural adaptation cannot be excluded (Extended Data Fig. 2). The antagonistic relationship of Lhca9^2^ and Lhca2, and the assembly state of PsaH might further reflect a regulation of PSI dimerization (Fig. 3a).

**Fig. 3:**
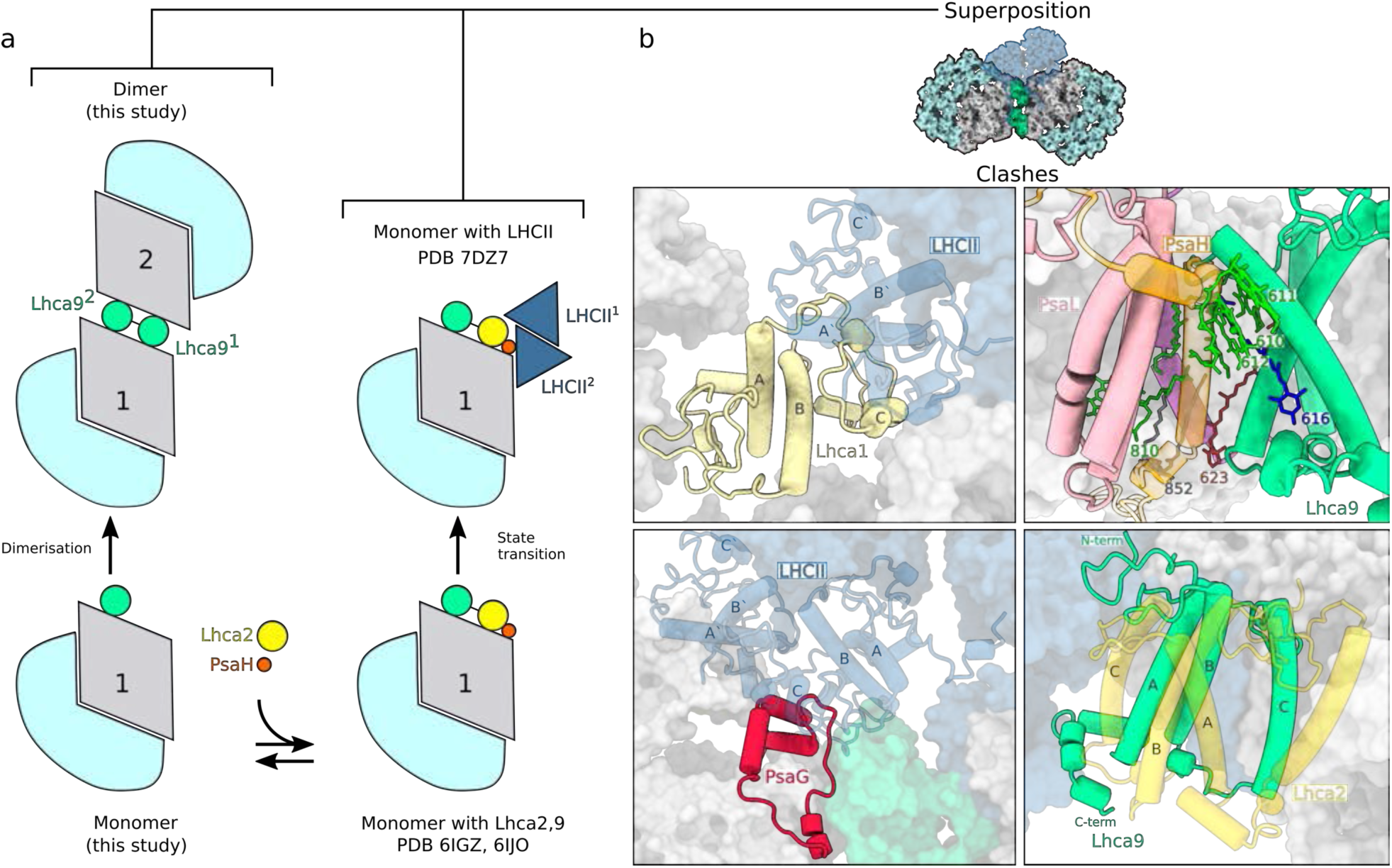
PSI dimer and LHCII association in *C. reinhardtii*. **a,** Schematic view of PSI divarication. The pathway towards state transition or dimer is dependent on presence/absence of PsaH and Lhca2. **b,** Superposition of PSI dimer with PSI-LHCII shows that state transition (PDB ID: 7DZ8, transparent) would result in clashes of Lhca1 (top left) and PsaG (bottom left) with LHCII. PsaH (top right), and Lhca2 (bottom right) would clash with the Lhca9 from the neighbouring PSI monomer.

PsaH is a 11-kDa transmembrane protein that is imported into chloroplasts and peripherally associates with the PSI reaction centre on the opposite side to the LHCI belt ^27^. A recent study of PSI biogenesis apparatus showed that PsaH is assembled in a separate protein module that comes before Lhca2 assembly ^28^. *Arabidopsis* mutants of PsaH can grow photoautotrophically ^29^, and structures of functional algal PSI particles that lack PsaH have been determined ^2, 14^. In addition, RNA-Seq analysis in algae further confirmed that expression of PsaH is regulated upon physiological stimuli ^30^. It is also of note that PsaH, unlike other PSI subunits, was found to be specifically enriched within the pyrenoid tubules ^31^. Together, these data suggest that PsaH- lacking complexes represent a previously overlooked functional form of PSI, where PsaH is either downregulated or could not be assembled. Building on these data, our structure of PSI-LHCI dimer further shows that PsaH/Lhca2 lacking particles can associate with each other to form larger complexes (Fig. 4). This feature is likely to be conserved in plants, as larger than monomer fractions of a plant PSI have been reported ^32^, and 2D projections from negative stain images of *Arabidopsis* PSI suggest the presence of a putative Lhca1/Lhca4 heterodimer at the PsaL pole that is analogous to alga ^33^. The dense organization would be beneficial for membrane crowding and compartmentalization of PSI, even if the two monomers are not energetically coupled.

**Fig. 4:**
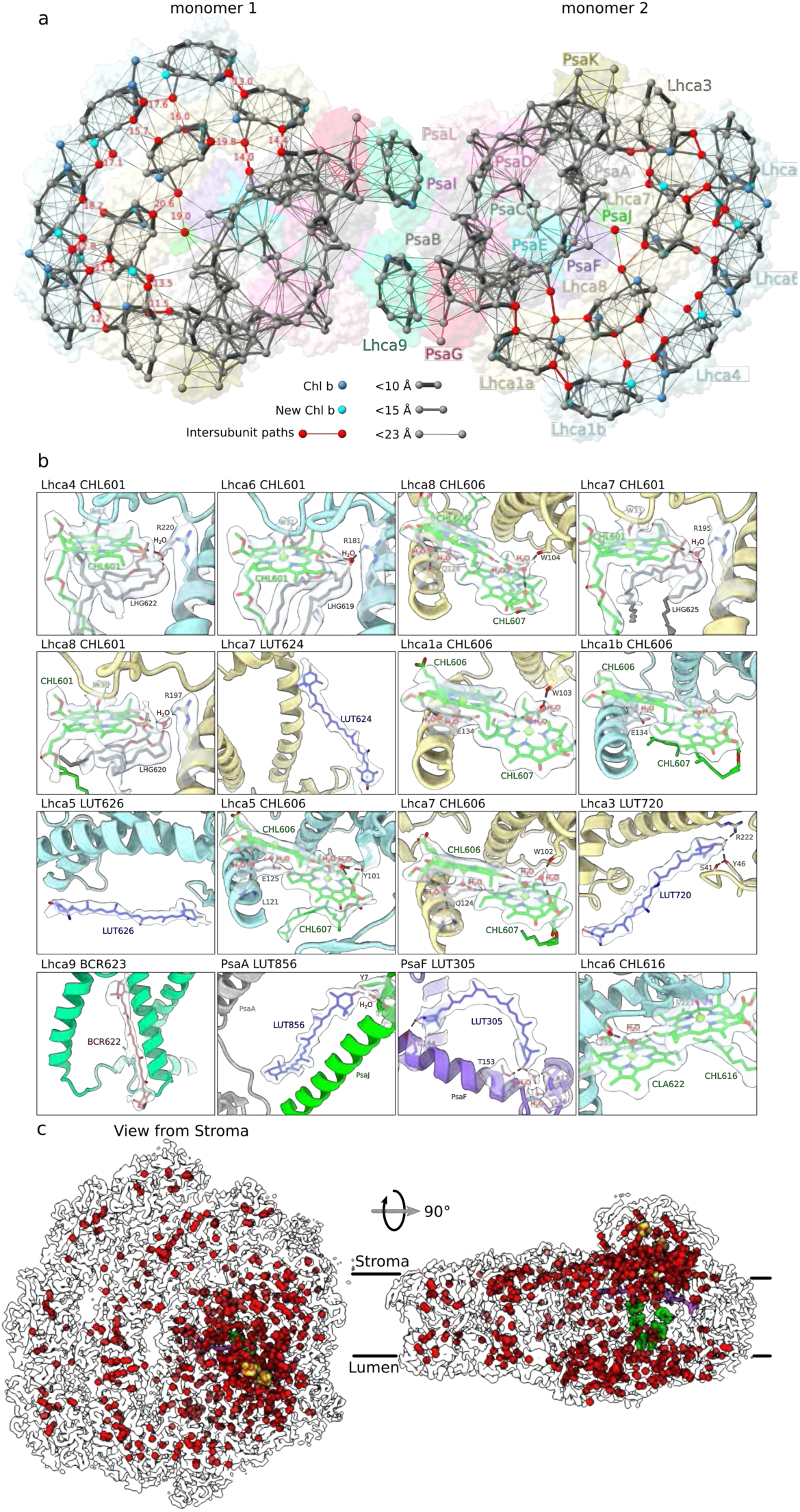
High resolution features of co-factors and solvation. **a**, Energy transfer pathways. Distribution of chlorophylls indicated by Mg atoms. Chl *a* gray, Chl *b* blue, newly identified Chl *b* cyan, pathways within 23 Å are connected by lines, and the line width reflects the distance. The most likely intersubunit pathways are red. **b**, Close up views of newly identified elements, coordination and chemical environment with the density map. **c**, Water molecules modeled in PSI are shown as red spheres (oxygen atoms) in the outline of the map, chlorophylls green, phylloquinones purple, iron-sulphur clusters yellow-red.

For other model organisms, such as *Saccharomyces cerevisiae*, a whole-cell modeling revealed that metabolic strategies are driven by specific protein expression profiles and compartment-specific proteome constraints ^34^. This allows eukaryotic cells to adapt metabolically to nutrient and proteome limitations. Crowding of large complexes in the membrane systems of thylakoid ^35, 36^, mitochondrial cristae ^37^, as well as in the cytosol, e.g. ribosome dimers ^38^ is an established and conserved mechanism for a more efficient differentiation of functions, ecological adaptation, and material storage during less active periods. Since the basic structure of the PSI core is rigid and it operates according to conserved mechanisms, the regulation by PsaH/Lhca2 provides a degree of flexibility on the macromolecular level that would allow the photosynthetic apparatus to adapt to stresses and tolerate changes in the range of light intensities that might involve membrane reorganisation ^15, 39^. Thus, PsaH/Lhca2 appears to be a regulator that defines a late assembly pathway and coordinates the macromolecular organization of PSI in chloroplasts (Fig. 3, Extended Fig. 8). This regulatory role could also be mediated through post-translational modifications such as phosphorylation in *C. reinhardtii* ^40^ or acetylation in *Arabidopsis* ^41^. In *C. reinhardtii*, a consequence of PsaH regulation would be the differential binding of Lhca2. Reduction of Lhca2 alters regulation of photosynthetic electron transfer and hydrogen production, suggesting further potential functional consequences of membrane reorganisation and PSI-remodelling ^26^.

### Coupling of PSI monomers in the dimer

To further extrapolate potential conformational changes during the dimerization of PSI, we applied multi-body refinement analysis of the PSI dimer using the two monomers as bodies (Extended Data Fig. 9). The analysis indicated no distinct conformational states, but instead revealed continuous motions in the three eigenvectors describing a relative movement of the monomers in relation to each other (Extended Data Fig 9a). The intrinsic flexibility is dominated by combinations of all three rotations of one monomer with respect to the other up to 13° (Extended Data Fig. 9b-d). Therefore, excitation energy transfer between the PSI monomers in the dimeric scaffold would also depend on degrees of rotation around the identified pivot points. Specifically, three chlorophylls are found within a potential cross-monomer excitation-sharing: CLA807 (PsaB), CLA604 (Lhca9^2^), CHL606 (Lhca9^2^), and the distance between them is ∼20 Å in the consensus map (Fig. 2). While such a positioning might suggest direct coupling, the multi-body analysis indicates considerable variability (Extended Data Fig. 9e,f). Therefore, similar to the cyanobacterial PSI dimer, an excitation coupling between the two monomers is less favourable *in vitro*, and this is consistent with measurements of 77 K fluorescence spectra that showed only a minor shift between monomer and dimer (Extended Data Fig. 1). However, *in vivo* the observed PSI-LHCI dimer conformation, and therefore the distance between the chlorophylls at the interface, could also be affected by a local membrane curvature.

### High resolution features and solvation of PSI

In our PSI monomer reconstruction, the resolution in the core is ∼2.1 Å, and in the LHCI inner belt 2.1-2.5 Å, revealing unprecedented structural details of chlorophylls, carotenoids, and 621 water molecules (Fig. 4, Supplementary Table 2). The map can improve the level of detail not only when compared to the previous cryo-EM studies of algal PSI ^1–3, 17^, but also the plant PSI maps obtained by X-ray crystallography ^19, 20^ (Extended Data Fig. 10). The quality of the data aided in improving the previous models in functionally important regions. This includes the identification of nine Chl *b* molecules, two newly modelled luteins, a beta-carotene and more accurate estimation of the coordination of 53 chlorophylls (Fig. 4b, Extended Data Fig. 11). Particularly, Chl *b* molecules are identified at positions 601 and 606 in Lhca4, Lhca5, Lhca6, Lhca7 and Lhca8. The two newly modeled luteins 720 and 626 are in the N-terminus of Lhca3 next to Chl *a* 614, and in Lhca5 next to Chl a 617, respectively (Fig. 4b). The newly modeled beta-carotene 622 is in Lhca9 and could be identified due to structural stabilization of the interface region in the dimer (Fig. 4b). Since chl *b* limits free diffusion of excitation energy ^42^, some of the new assignments affect the energy pathways between the antenna proteins. Together with the new structural data, this allowed us to produce a more accurate map of the energy channelling in PSI based on the new model (Fig. 4a).

Another striking feature of the high resolution cryo-EM map is resolvable density for multiple newly detected water molecules, which particularly aided in modeling the coordination of chlorophylls (Fig. 4c, Supplementary movie 1). Thus, we report the most complete available experimental picture of a chemical environment for chlorophyll binding (Supplementary Table 3). Particularly, it allows distinguishing between mono- and di-hydrated forms, which largely escaped detection by X-ray crystallography (Extended Data Fig. 12). This is mechanistically important because the di-hydrated derivative is chemically more stable, as illustrated by quantum chemical calculations ^43^. We observe that, other than the previously reported CLA824 ^20^, only two waters can be involved in penta-coordinated Mg for all the chlorophylls. Remarkably, waters play a coordinative role for most of Chl *b*, for which the relative ratio of water coordination is four times higher than for Chl *a* (Supplementary Table 3). The difference between Chl *a* and Chl *b* is a methyl versus a formyl group, thus water serves as a hydrogen bond donor to the latter, while it also interacts with charged/polar protein residues or lipids. Therefore, the immediate surrounding of Chl *b* molecules is more enriched with non-protein material than previously thought, which plays a role in tuning the photophysics and the transport properties of excitation energy in PSI. Together, the presented model now allows for comparison of PSI phylogenetic conservation also on the level of chlorophyll coordination and solvent positioning.

### Structure of PSI:Pc complex

On the lumenal side of PSI, we observed a density corresponding to the associated electron donor Pc, which binding has been stabilised by crosslinking (see Methods). The bound Pc is found on PSI monomers and dimers. Signal subtraction, followed by focused 3D classification, allowed us to rigid body fit a model for Pc into the density at the local resolution of ∼3.5 Å (Extended Data Fig. 3). We then performed flexible fitting using self-similarity-restraints in *Coot* ^44, 45^. With respect to the PSI-Pc interactions, comparison between our model with a plant counterpart ^46, 47^ revealed two main differences (Fig. 5). In *C. reinhardtii*, the negatively charged residues of the Pc acidic patch are shifted by ∼5 Å due to the missing residues P58-E59, and therefore, the interaction with K101 of PsaF is weakened (Fig. 5a,b). Instead, the binding strength is compensated with the PsaF region K78-K92, which has six lysines (78, 81, 82, 85, 89, 92) increasing a positively charged concentration at a distal site (Fig 5c-e), thus supporting additional electrostatic interactions with the acidic residues of Pc. The importance of the distal lysine residues is supported by site-directed mutagenesis of the *C. reinhardtii* N-terminal PsaF domain and functional analyses of electron transfer between Pc and mutant as well as wild type PSI ^48^. Thus, algal and plant Pc have adapted their slightly different interfaces for optimal interactions with PSI.

**Fig. 5:**
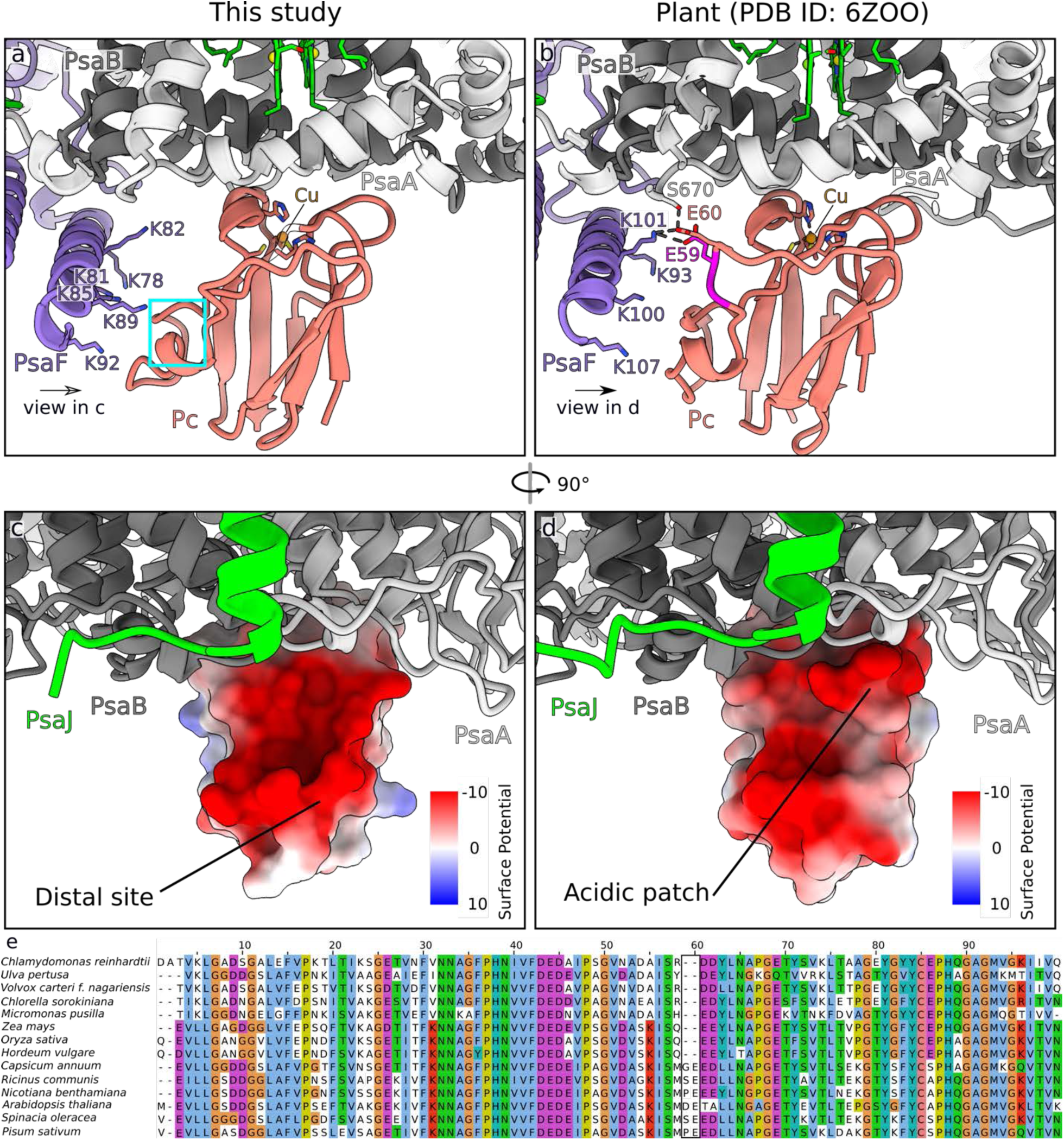
Plastocyanin binding site. **a**, Pc binding in *C. reinhartdii* (current work). The positively charged residues of PsaF stabilize the interactions. The corresponding Pc region that deviates from the plant counterpart is cyan. **b**, Pc binding in plants ^29^. The two inserted residues are magenta. **c**, 90° rotated view with Pc surface shown with Coulomb potential from the interface. **d**, the same view for a plant counterpart (PDB ID: 6ZOO), illustrating that the acidic patch is shifted. **e**, Multiple sequence alignment of Pc from different species of the green lineage (algae and plants) showing that the inserted residues 58, 59 occur in a subset of plants and do not represent a general case.

In summary, this study illustrates that PSI-LHCI can dimerise and explains how the process is structurally regulated by subunits PsaH, Lhca2, and Lhca9. It remains to be seen how thylakoid regulatory networks manage to implement PsaH allocation strategies. The data provides the most complete description so far of the structure of PSI, including newly identified cofactors that play specific roles in accommodating the light harvesting and excitation transfer functions, as well as water molecules involved in chlorophyll coordination. Finally, the binding of electron donor Pc is resolved. Together, the data explain how the PSI is modulated to perform its functional and structural roles in a chloroplast.

## Methods

### Strains and growth conditions

Experiments were performed using a previously described strain expressing PsaB fused with His20-tag after the third residue from the N-terminus ^2^ and an *Δlhca9* insertional mutant ^49^ transformed with the corresponding PsaB His tag plasmid. Chloroplast transformation and transformant selection was performed as described previously. Further experiments were performed with the *C. reinhardtii* wild type strain 137c as well as *Δlhca2* insertional mutant ^45^ back-crossed into wild type 137c.

All strains were maintained on TAP medium, solidified with 1.5% w/v agar at 25 °C in the presence of ∼20 μmol photons m^-2^ s^-1^ photosynthetically active, continuous illumination. For experiments, strains were cultured in TAP medium on a rotary shaker (120 rpm) at 25 °C in the presence of ∼20 μmol photons m^-2^ s^-1^ photosynthetically active, continuous illumination.

### Purification of PSI

*C. reinhardtii* cells were incubated in anoxic conditions (∼10^8^ cells mL^-1^ in TAP medium + 10 mM glucose, 40 U mL^-1^ glucose oxidase, 50 U mL^-1^ catalase) and dim light (∼20 µmol photons m^-2^ s^-1^) for 60 min. All following steps were performed at 4 °C and dim light. Cells were disrupted in 0.33 M sucrose, 25 mM HEPES-KOH pH = 7.5, 5 mM MgCl_2_, 1 mM PMSF, 1 mM benzamidine, 5 mM aminocaproic acid with a nebulizer (2 bar, two passages). Broken cells were centrifuged at 20,000 rpm for 10 min (Beckman Coulter JA-25.50 rotor). The pellet was carefully resuspended in 0.5 M sucrose, 5 mM HEPES-KOH pH = 7.5, 10 mM EDTA, 1 mM benzamidine, 5 mM aminocaproic acid with a potter homogenizer. The resuspended material was layered on top of a sucrose density step gradient (1.8 M and 1.3 M sucrose, 5 mM HEPES-KOH pH = 7.5, 10 mM EDTA, 1 mM benzamidine, 5 mM aminocaproic acid). Thylakoid membranes were extracted via ultracentrifugation at 24,000 rpm for 1 h and 20 min (Beckman Coulter SW 32 Ti rotor). Thylakoids were collected from the step gradient interphases with a Pasteur pipet, diluted four times with 5 mM HEPES-KOH pH = 7.5 and centrifuged at 21,5000 rpm for 20 min (Beckman Coulter JA 25.50).

Isolated thylakoids were set to 1 mg chl mL^-1^ in 5 mM HEPES-KOH pH = 7.5 and solubilized by addition of an equal volume of 2 % α-DDM for 10 min. Unsolubilized material was separated by centrifugation. The supernatant was diluted four times to 125 µg chl mL^-1^ and 0.25 % α-DDM. The sample was loaded onto a TALON metal affinity column (1 mL resin mg chl^-1^) in 5 mM HEPES-KOH pH = 7.5, 100 mM NaCl, 5 mM MgSO4, 10 % glycerol at a flow rate of ∼0.5 mL min^-1^. The column was washed with 10 times the bed volume of 5 mM HEPES-KOH pH = 7.5, 100 mM NaCl, 5 mM MgSO_4_, 10 % glycerol, 0.02 % α-DDM at a flow rate of ∼1 mL min^-1^. A second wash was performed with 10 times the bed volume of 5 mM HEPES-KOH pH = 7.5, 100 mM NaCl, 5 mM MgSO_4_, 10 % glycerol, 0.02 % α-DDM and 5 mM imidazole at a flow rate of ∼1 mL min^-1^. The PSI was eluted with 5 mM HEPES-KOH pH = 7.5, 100 mM NaCl, 5 mM MgSO_4_, 10 % glycerol, 0.02 % α-DDM and 150 mM imidazole. The PSI was concentrated with a spin column (regenerated cellulose: 100,000 MWCO) to ∼ 1.5 mg chl mL^-1^, diluted five times with 30 mM HEPES-KOH pH = 7.5, 0.02 % α-DDM and reconcentrated twice.

PSI-Pc crosslinking was performed in 30 mM HEPES-KOH pH = 7.5, 1 mM ascorbate, 0.1 mM diaminodurene (a redox mediator between ascorbate and Pc), 3 mM MgCl_2_ with 0.1 mg chl mL^-1^ PSI particles and 20 µM activated Pc for 45 min at room temperature. Pc activation was performed in 10 mM MOPS-KOH pH = 6.5 with 100 µM recombinant Pc, 5 mM EDC and 10 mM N-hydroxysuccinimide for 20 min at room temperature. The crosslinker was removed and the buffer exchanged to 30 mM HEPES-KOH pH = 7.5 via a PD G25 desalting column followed by ultrafiltration with a centricon (regenerated cellulose: 10,000 MWCO).

The crosslinked PSI-Pc particles (∼ 60 µg chl per gradient) or solubilized thylakoids of non-tagged strains (∼ 250 µg chl per gradient) were loaded onto a 1.3 M – 0.1 M sucrose density gradient including 5 mM HEPES-KOH pH = 7.5 and 0.02 % α-DDM. PSI fractions were collected after ultracentrifugation at 33,000 rpm (Beckman Coulter SW 41 Ti) for 14 h (Extended Data Fig. 1b and 6). Prior to further analysis, sucrose was removed via a PD G25 desalting column followed by concentration with a spin column (regenerated cellulose: 100,000 MWCO).

### Biochemical analysis of PSI

For SDS-PAGE (Extended Data Fig. 1c-e), samples were adjusted to 1 µg chl, supplemented with loading buffer and incubated at 65 °C for 15 min. Proteins were separated by 13% (w/v) SDS-PAGE ^50^. Gels were stained with Coomassie Brilliant Blue R-250 or blotted onto nitrocellulose membranes (Amersham) and detected by specific primary antibodies: PsaF ^51^, PC (Agrisera), PsaA, Lhca5, Lhca2, Lhca9 ^4^, Lhca3 ^52^, PsaD ^53^, PsaG and Lhcb/a ^54^. The antibody against PsaA was raised using the following peptides STPEREAKKVKIAVDR and VKIAVDRNPVETSFEK and was obtained from Eurogentec (Belgium). Secondary antibodies for ECL detection were anti-rabbit (Invitrogen).

For quantitative analysis by mass spectrometry, removal of sucrose and protein digestion was carried out following the FASP protocol, using 2 µg of sequencing grade trypsin (Promega) per fraction ^55^. Iodoacetamide and dithiothreitol used in the original protocol were replaced by chloroacetamide and tris (2-carboxyethyl) phosphine, respectively. After over-night digestion at 37°C, samples were acidified by adding trifluoroacetic acid (TFA) to a final volume of 0.1%. Five percent of the peptide solution were desalted using self-made Stage tips according to established protocols ^56^. Desalted peptides were dried by vacuum centrifugation and stored at - 20°C until further use. The LC-MS/MS system consisted of an Ultimate 3000 RSLC nanoLC System (Thermo Fisher Scientific) coupled via a Nanospray Flex ion source (Thermo Fisher Scientific) to a Q Exactive Plus mass spectrometer (Thermo Fisher Scientific). Samples were reconstituted in 5 µl of 2 % (v/v) acetonitrile/0.05% (v/v) TFA in ultrapure water (eluent A1), loaded on a trap column (C18 PepMap 100, 300 µM x 5 mm, 5 µm particle size, 100 Å pore size; Thermo Fisher Scientific) and desalted for 3 min at a flow rate of 15 µL min^-1^ using eluent A1. Subsequently, the trap column was switched in-line with an Acclaim PepMap100 reversed phase column (75 µm x 50 cm, 2 µm particle sizes, 100 Å pore size; Thermo Fisher Scientific) for peptide separation. The mobile phases were composed of 0.1 % (v/v) formic acid in ultrapure water (eluent A2) and 80 % (v/v) acetonitrile/0.08 % (v/v) formic acid in ultrapure water (B). Flow rate was 250 nL/min. The following gradient was applied: 5-35 % B over 105 min, 35-99 % B over 5 min, 99 % B for 20 min. MS full scans (scan range m/z: 350–1400, resolution 70,000 at m/z 200, AGC target value 3e6, maximum injection time 50 ms) were acquired in data-dependent mode, dynamically selecting the 12 most abundant precursor ions for fragmentation by higher-energy C-trap dissociation (HCD, 27 % normalized collision energy, resolution 17,500 at m/z 200, precursor isolation window 1.5 m/z). Dynamic exclusion was set to ‘auto’ (chromatographic peak width: 15 s). AGC target value and intensity threshold for MS/MS were 5e4 and 1e4, respectively, at 80 ms maximum ion fill time. Singly charged ions, ions with charge state 5 or above and ions with unassigned charge states were rejected. Internal lock mass calibration was enabled on m/z 445.12003. LC-MS/MS data was processed in MaxQuant 1.6.14 for protein identification and label-free quantification ^57^. Default settings were used, except for calculation of Intensity-Based Absolute Quantitation (iBAQ) values and “second peptide search”, which were enabled and disabled, respectively. iBAQ values were normalized to the PSI core subunit PsaB and values below 21 were excluded as they represent already low intensity values, which might not be reliable. Spectra were searched against a concatenated database containing protein sequences based on the *Chlamydomonas* v5.6 gene models (Joint Genome Institute, www.phytozome.org), as well as sequences of chloroplast- and mitochondrial-encoded proteins (GenBank BK000554.2 and NC_001638.1). Carbamidomethylation of cysteines was set as a fixed modification. Oxidation of methionine and acetylation of protein N-termini were considered as variable modifications. A false discovery rate (FDR) of 1% was applied to peptide and protein identifications. LFQ data was imported into Perseus (version 1.6.15.0) ^58^ for log2- transformation, and contaminants, proteins only identified by site and reverse hits were removed. Room temperature absorption spectra (300 – 750 nm) of PSI monomer and dimer fractions (Extended Data Fig. 1g) were measured with a UV-/Vis spectrophotometer (V-650, Jasco) at 10 µg chl mL^-1^ and normalized to the red region. Fluorescence emission spectra (650 – 780 nm) of PSI fractions (Extended Data Fig. 1h) were recorded at 77 K at 1 µg chl mL^-1^ with a spectrofluorometer (P-6500, Jasco) upon excitation at 436 nm. Spectra were normalized to the maximum of the emission peaks and smoothed according to ref 59. For 2D-PAGE, thylakoids (0.8 mg chl mL^-1^) isolated from control and anoxic 137c wild type cells were solubilized with 0.9 % β-DDM for 20 min. 2D-PAGE and silver staining ^52^ (Hippler et al., 2001) were performed as described. Excised spots from silver-stained blue native PAGE gels were destained by incubation with 15 mM potassium hexacyanoferrate (III)/50 mM sodium thiosulfate for 8 min, and then submitted to tryptic in-gel digestion as described ^60^. No reduction and alkylation of cysteines was performed. LC-MS/MS was implemented as described ^53^, where an Ultimate 3000 nano-LC system was coupled via a nanospray source to an LCQ Deca XP Plus mass spectrometer (Thermo Finnigan).

LC-MS/MS data were processed with Proteome Discoverer (Thermo Fisher Scientific, version 2.4). Raw files were searched using the SequestHT and MS Amanda algorithms against a concatenated database containing sequences of nuclear- (Chlamydomonas v5.6 gene models, www.phytozome.org), chloroplast- (GenBank BK000554.2) and mitochondrial-encoded (NC_001638.1) proteins Search settings were: precursor and fragment mass tolerances 250 ppm and 0.25 Da, respectively; minimum peptide length: 6; maximum of missed cleavages: 2; variable modifications: oxidation of methionine, N-acetylation of protein N-termini. Identifications were filtered to achieve a peptide and protein level false discovery rate of 0.01.

### Cryo-EM data collection and processing

3 μL of purified PSI complex at 1 mg Chl mL^−1^ were applied to glow-discharged (GloQube Quorum, 40 sec, 20mA) holey carbon grids (Quantifoil 300 Au R1.2/R1.3, Electron Microscopy Sciences) and vitrified using a Vitrobot MKIV (2 sec blotting time, 4°C, 100% humidity). The data collection was carried out using a 300KV Titan Krios G2 Microscope (Thermo Fisher Scientific) equipped with a Gatan Bioquantum energy filter and a K3 Summit direct electron detector (Ametek). Movies were recorded using counting mode at a magnification of ×105,000 corresponding to a calibrated pixel size of 0.84 Å. The dose rate was 15.27 e/px/sec and the exposure time was 3 sec divided into 45 frames leading to a total dose of 45.8 e Å^-2^. EPU was used to collect 17,439 movies with a defocus range from -0.7 to -2.5µm. Data statistics are shown in Supplementary Table 2.

Extended Data Fig. 3 shows the processing scheme applied. The pre-processing steps were performed using cryoSPARC 3.1.0 ^61^. Movie stacks were motion corrected and dose weighted using MotionCor2 ^62^. Contrast Transfer Function (CTF) of the motion corrected micrographs was estimated using CTFFIND4 ^63^. Blob picker and then template picker were used to pick 440,494 particles and 2D classification in cryoSPARC was performed. Dimeric particles were separated from monomeric by inspection of the 2D-class averages and for each sub-population an *Ab initio* model was generated using cryoSPARC applying C2 and C1 symmetry, respectively. For each model homogenous refinement was performed leading to a nominal resolution of 3.7 Å for the dimer and 3.0 Å for the monomer. Particles (dimer: 69,144 monomer: 123,746) were converted into a Star file format ^64^ and were imported into RELION 3.1.beta ^65, 66^. Particles were re-extracted (un-binned) and processed in RELION using a box size of 700 pixel and 500 pixel for the dimer and monomer, respectively. 3D Refinement followed by 3D classification was performed imposing C2 symmetry for the dimer and C1 for the monomer. A subset of high quality particles was selected for the dimer and monomer and subjected to 3D refinement which resulted in a resolution of 3.3 Å for the dimer and 2.9 Å for the monomer. CTF refinement ^67, 68^ followed by 3D refinement and Bayesian polishing followed by another round of CTF refinement was performed for the dimer as well as for the monomer. A final 3D refinement resulted in an overall resolution of 2.97 Å for the dimer and 2.31 Å for the monomer. The resolution of the dimer could be further improved to 2.74 Å by using signal subtraction of one monomer followed by symmetry expansion and 3D refinement applying C1 symmetry.

In order to increase the number of particles for classification on the Pc region, Dataset 2 was collected from the same dimer band, but with a pixel size of 0.51 Å. The dataset was processed with cryoSPARC 3.1.0 ^61^. After template picking 864,399 particles were extracted. With a small subset of the extracted particles an *Ab initio* reconstruction was generated followed by Heterogeneous Refinement using 5 classes, one of which contained *Ab initio* reconstruction as reference. The class containing the PSI monomer was then subjected to Homogeneous Refinement in cryoSPARC 3.1.0 resulting in a reconstruction at 3.88 Å resolution. The particles were then exported to RELION ^65^ and 3D classification was performed. The class that contained 88,219 good particles was used for further refinement which improved the overall resolution to 3.5 Å. Applying CTF-refinement and Bayesian Polishing resulted in further improvement, and the final nominal resolution is 2.68 Å. The data was then merged with the monomer (Dataset 1) and signal subtraction followed by focused classification using a mask around the Pc region was performed. A class with 66,080 particles showed the best density for Pc which was used for model building. The workflow is further illustrated in the Extended Data Fig 3a.

For analysis of the motion between the two monomers of the dimer we performed a Multibody Refinement in RELION ^69^ followed by a principal component analysis using the program relion_flex_analyse. Two bodies were chosen, one for each monomer, resulting in twelve Eigenvectors describing the motion. Ten maps for each of the three Eigenvectors which describe about 78% of the motion in the data were printed out and the maps with the extreme positions (map1 and 10) were used to fit the models that are shown in Extended Data Figure 8. A Python script was used to estimate the distances between the chlorophylls at the dimer interface for each particle in the data and to plot the results as histograms as depicted in Extended Data Figure 9e,f.

### Model building and refinement

Initially, the available model of the PSI structure (PDB ID: 6JO5) of *C. reinhardtii* was rigid body fitted into the 2.74 Å map of the symmetry expanded dimer using Chimera v 1.14 ^70^. Model building and real-space refinement was then carried out using *Coot* v9.1.4 ^44, 45^ to complete one monomer. Two copies of the completed monomeric model were then rigid body fitted into the C2 generated 2.97 Å dimer map using Chimera. The model of the monomer was then fitted separately in the highest resolution 2.3 Å map of the monomer. All protein residues as well as pigments were fitted using *Coot* ^44, 45^ with locally optimized map weights. The cis-trans isomerism of each pigment was judged based on density and modeled accordingly. Newly identified chlorophylls and carotenoids were modeled, when the experimental evidence (density map) supported and the chemical environment matched the surrounding of the pigments. For carotenoids, the density that clearly showed the characteristics of an elongated tetraterpenoid with densities for the four methyl groups, two sticking out on each side of the chain, was identified as a new carotenoid binding site. To further analyse the identity of the corresponding pigment possible candidates were fitted and compared. A carotenoid that fitted best in terms of density and chemical environment was then selected. In case of luteins, the oxygen of the cyclohexane- ring was the main criterion for pigment identity because it is involved in hydrogen bonding. For the Chl *b* identification the densities for the aldehyde group needed to be present as well as the hydrogen bonding occurring with a water molecule that are usually stabilized by other chlorophylls, lipids, and protein side chains. Water molecules involved in pigment interactions were placed manually. All other water molecules were picked by *Coot* ^44, 45^ with the auto picking function followed by manual inspection and correction. All high resolution features were modelled using *Coot* ^44, 45^ until the model was completed. For all modeling steps restraint files for pigments and ligands were used which were generated using the Grade server (http://grade.globalphasing.org). Restraint files were adopted manually if it was required.

For plastocyanin, a model was generated using SWISS model ^71^. The model was then rigid body fitted using Chimera. Rotamers were corrected for the residues that were allowed due to the better local densities. Self-restraints in Coot were activated followed by flexible fitting into the density. All models were refined using Real-Space-Refine from the PHENIX suite ^72^ using the Grade server restraint files for the ligands and a distance .edit file which was generated by Ready-set in PHENIX. Further, hydrogen atoms were added for refinement to the model using Ready-set. The refinement protocol was optimized using different weight parameters. The refinement statistics are shown in Supplementary Table 2. Multiple rounds of validation and model building were carried out using MolProbity ^73^ and *Coot* ^44, 45^. For further validation the PDB Validation server was used (https://validate-rcsb-2.wwpdb.org).

The structure was analysed using *Coot* and Chimera. Figures were prepared using ChimeraX ^74^.

### Data availability

The atomic coordinates have been deposited in the Protein Data Bank (PDB) and are available under the accession codes: PDB ID: XXX for the dimer, PDB ID: XXXX for the symmetry expanded dimer, PDB ID: XXXX for the high resolution monomer, and PDB ID: XXXX for the PSI-Pc model. The local resolution filtered maps, half maps, masks and FSC-curves have been deposited in the Electron Microscopy Data Bank with the accession codes EMD-XXXXX for the dimer, EMD-XXXXX for the symmetry expanded dimer, EMD-XXXXX for the high resolution monomer, and EMD-XXXXX for the Plastocyanin map. Mass spectrometry datasets: Project Name: Intensity-based absolute quantification (iBAQ) of components of photosystem I monomers and dimers. Project accession: PXD026990. Project DOI: 10.6019/PXD026990. Project Name: Identification of photosystem I components from *Chlamydomonas reinhardtii* grown under oxic and anoxic conditions. Project accession: PXD027067.

## Acknowledgements

The cryo-EM data were collected at the SciLifeLab facility (funded by the KAW, EPS, and Kempe foundations) and the Karolinska Institutet 3D-EM facility. We thank D. Kimanius for providing a script for conversion of poses from Euler space into an orientation matrix in Cartesian space, S. Hawat for mass spectrometric analyses, V. Adlfar for isolation of Pc. Funding: Swedish Foundation for Strategic Research (ARC19-0051), Knut and Alice Wallenberg Foundation (2018.0080), Deutsche Forschungsgemeinschaft (739/13-2), Federal states (NRW 313-WO44A), German-Israeli Foundation for Scientific Research and Development (G-1483- 207/2018). AA is supported by the EMBO Young Investigator Programme, MH is supported by the RECTOR programme (University of Okayama, Japan).

## Author contributions

M.H. and A.A. designed the project. L.M., A.P.B. and A.N. prepared the sample for cryo-EM and performed initial screening. A.N. collected and processed the cryo-EM data and built the model. L.M., S.K., A.V.M. performed biochemical analysis. V.T. performed computational analysis. Y.T. provided antibodies. T.T.H.H. produced mutant strains. A.N. and A.A. analysed the structure and wrote the manuscript with contributions from L.M. and M.H. All authors contributed to the analysis and the final version of the manuscript.

---

**Extended Data Fig. 1:**
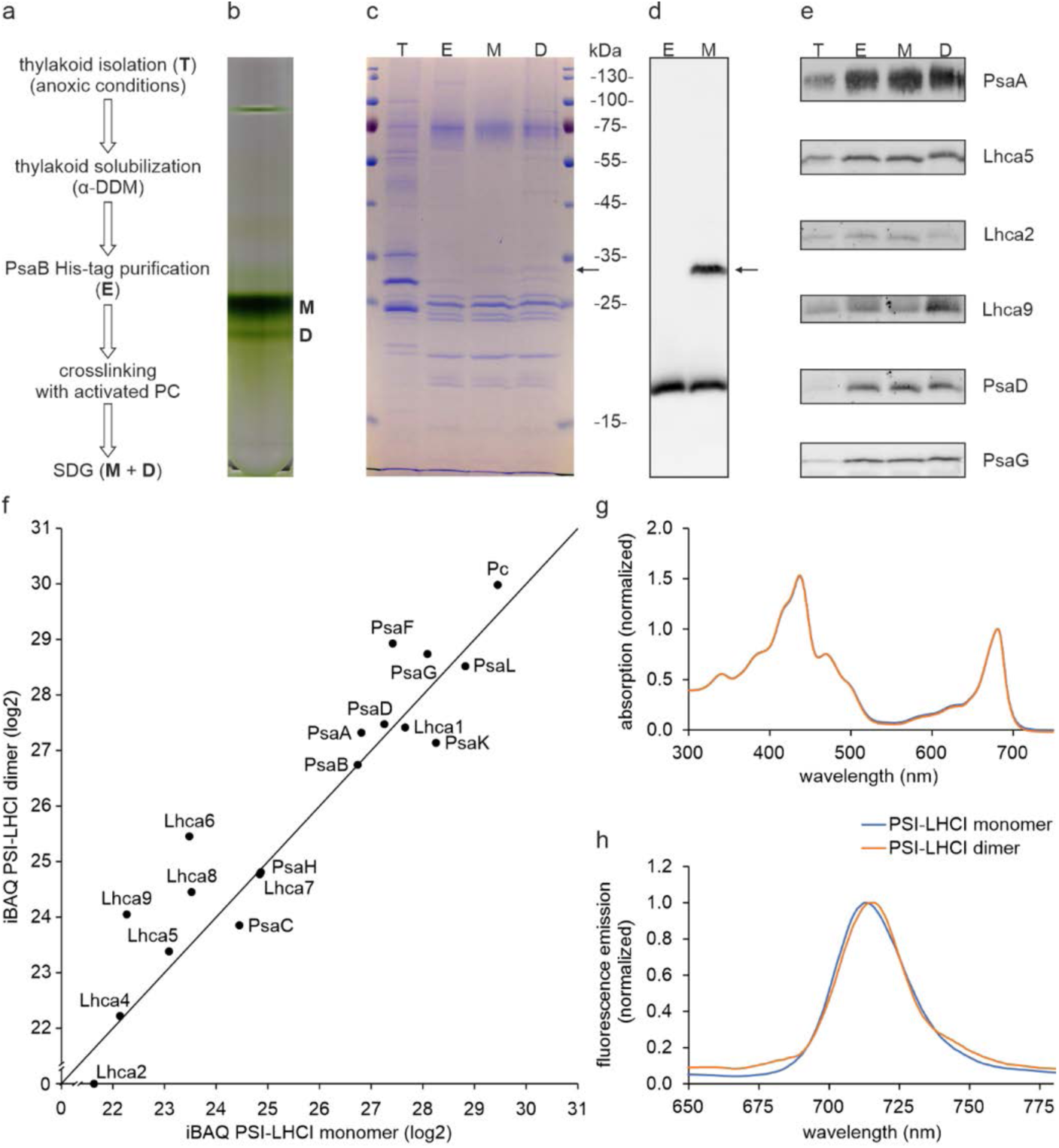
Sample preparation and characterisation. **a**, The experimental workflow. **b**, Sucrose density gradient affinity purified PSI (∼ 60 µg chl). Monomer (M) and dimer (D) fractions are indicated. **c**, SDS-PAGE analysis (1 µg chl). T: thylakoids; E: PSAB-His elution, M: PSI monomer, D: PSI dimer. The arrow indicates a putative PsaF-Pc crosslinked product. **d**, SDS-PAGE and Western Blot against PsaF confirming the cross-linking with Pc. **e**, SDS PAGE and Western Blot against PsaA, Lhca5, Lhca2, Lhca9, PsaD and PsaG confirming the presence of small PSI subunits as well as LHCI subunits from both sides of the PSI core. **f**, Quantitative mass spectrometry analysis of the PSI fractions from (**b**). iBAQ values are normalized to the PSI core subunit PsaB and values below 21 are excluded as they represent already low intensity values, which might not be reliable. PSI-LHCI subunits PsaE and Lhca3 were not detected at quantifiable intensities in this experiment. Detection of these small PSI and Lhca subunits via mass spectrometry depends on the presence of only a few proteotypic tryptic peptides, which could be missed, as measurements were done in a data dependent fashion (12 MS^2^ per MS^1^). **g**, Room temperature absorption spectra of PSI monomer and dimer fractions. The spectra are normalized to the red region. **h**, Low temperature (77 K) fluorescence emission spectra of PSI monomer and dimer fractions upon excitation of chlorophyll *a* at 436 nm. The spectra are normalized to the maximum of the emission peaks and smoothed according to ^42^. Notably, the presence of PsaH in the dimer sample, but its absence in the cryo-EM structure, indicates that PsaH is only loosely attached to the PSI dimer and readily lost during the cryo-EM analyses.

**Extended Data Fig. 2:**
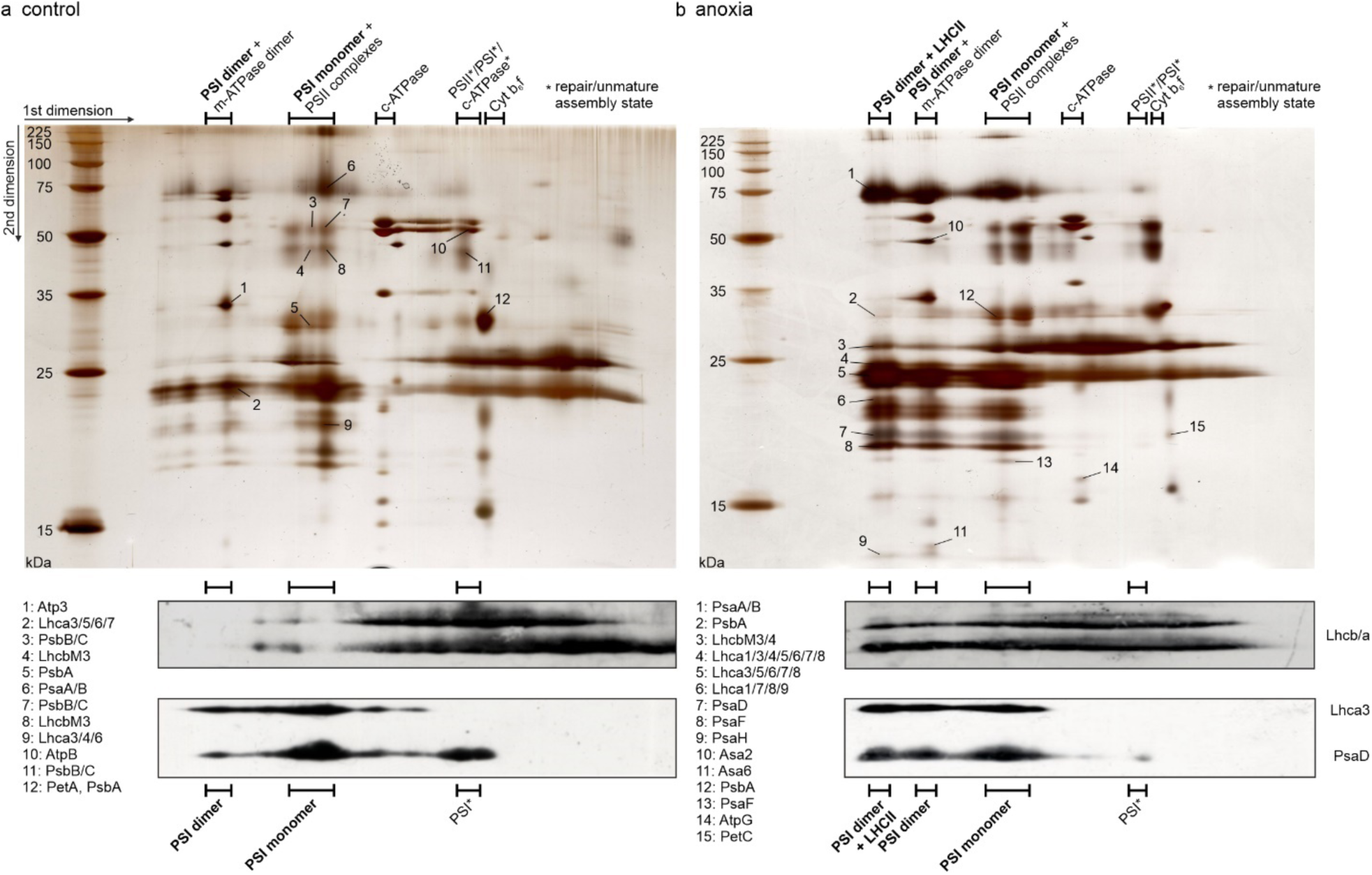
2D-PAGE of β-DDM solubilized *C. reinhardtii* wild type thylakoids. **a**, control conditions. **b**, low light and anoxic conditions. In the first dimension, multi-protein complexes were separated in their native form by blue native PAGE. In the second dimension, subunits of these protein complexes were separated by SDS-PAGE. “PSII complexes” refers to PSII complexes of different molecular weight due to varying extents of LHCII association. Labelled silver-stained spots were subjected to LC-MS/MS analysis. For convenience, the figure includes a condensed list of representative subunits. For a list of all proteins identified in the respective spots refer to Extended Data Table 1 and Supplemental Dataset 1. Western Blots of Lhcb/a, Lhca3 and PsaD indicate the distribution of LHCII, LHCI and the PSI core.

**Extended Data Fig. 3:**
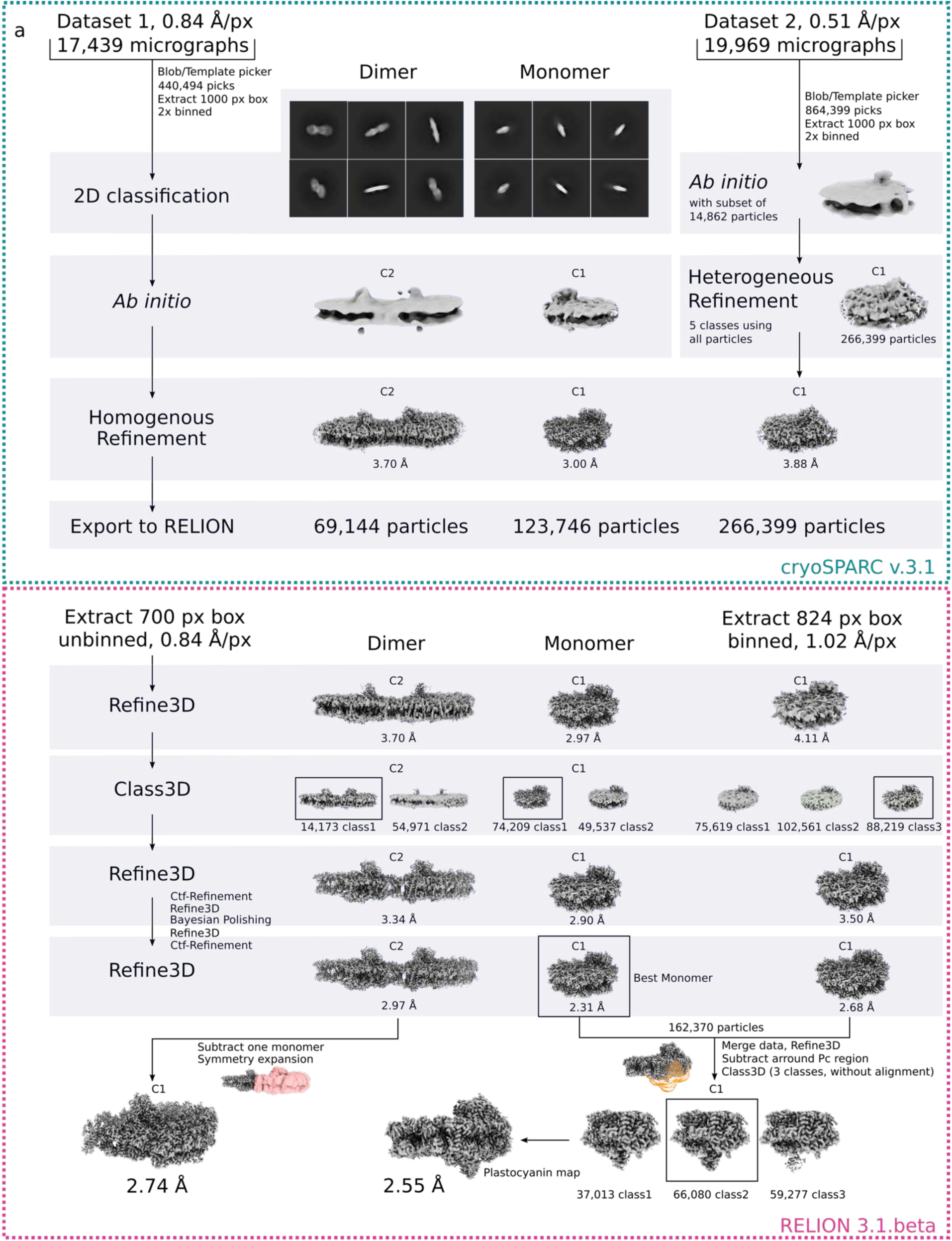
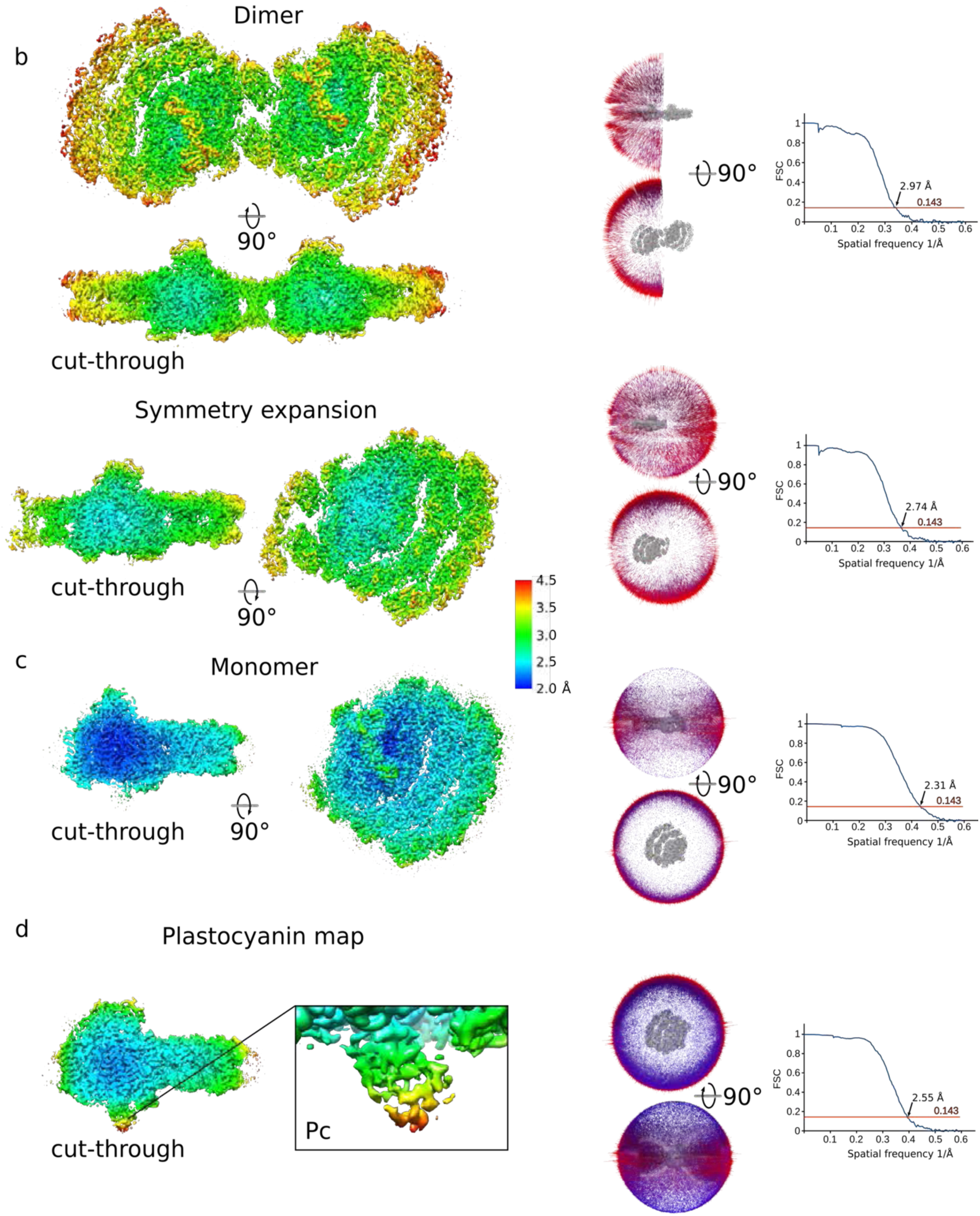
Cryo-EM processing workflow and local resolution. **a**, Flowchart of the data processing for the PSI dimer and monomer. The processing steps performed in cryoSPARC and RELION are indicated. To increase the numbers of Pc bound particles Dataset 2 was added for focused classification with signal subtraction. **b**, The map of the dimer and symmetry expanded dimer colored by local resolution in two orientations (left). Angular distribution of the dimer and symmetry expanded dimer and the FSC-curves are shown. **c**, The map of the monomer, angular distribution and FSC-curve. **d,** The map with the best density for plastocyanin, angular distribution and FSC-curve.

**Extended Data Fig. 4:**
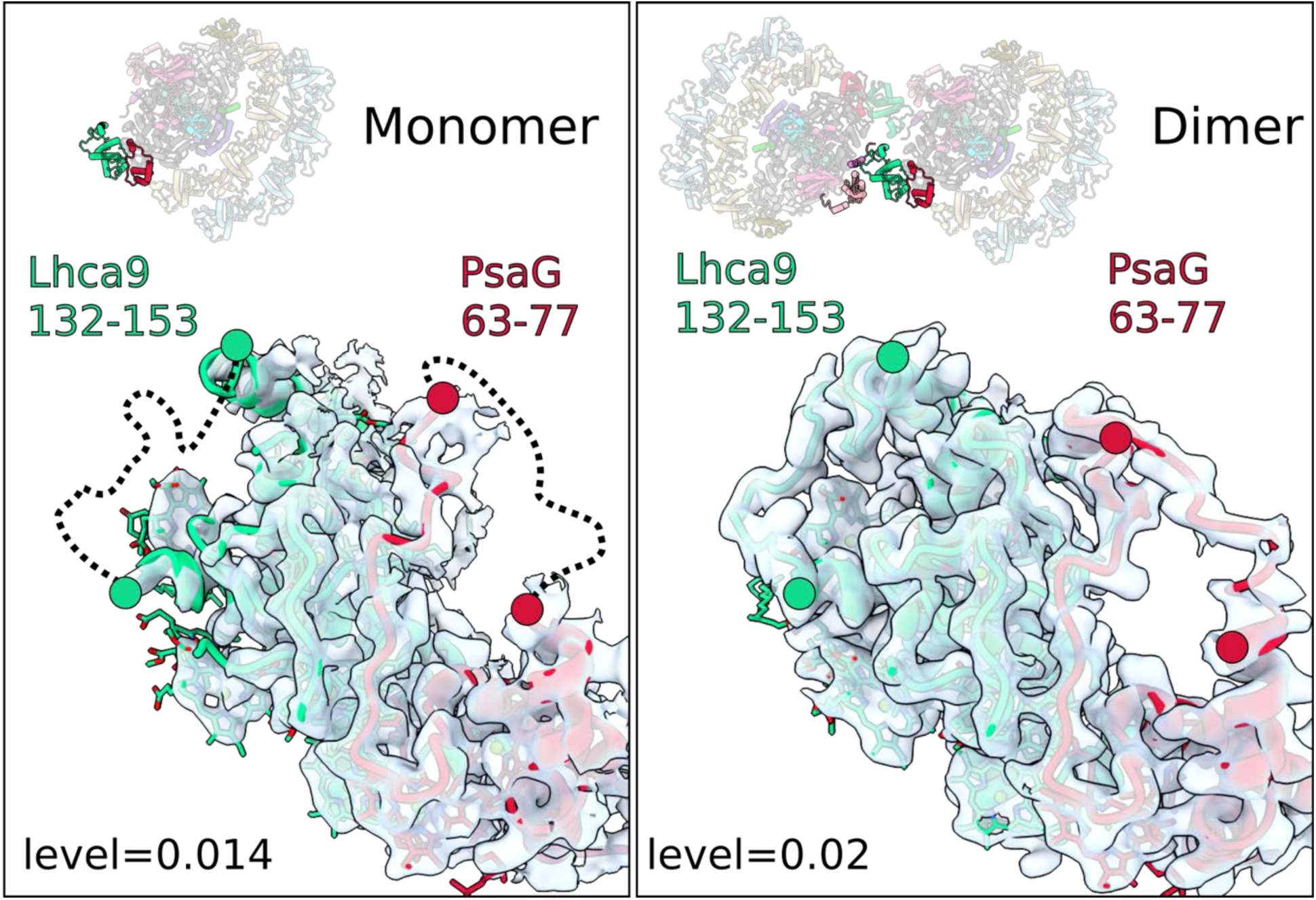
Stabilization of loops in Lhca9 and PsaG in PSI dimer. Left, in the monomer, the Lhca9 loop 132-153 and PsaG loop 63-77 are disordered (dashed lines). Right, in the dimer, both loop regions are ordered due to a stabilization by the neighbouring monomer.

**Extended Data Fig. 5:**
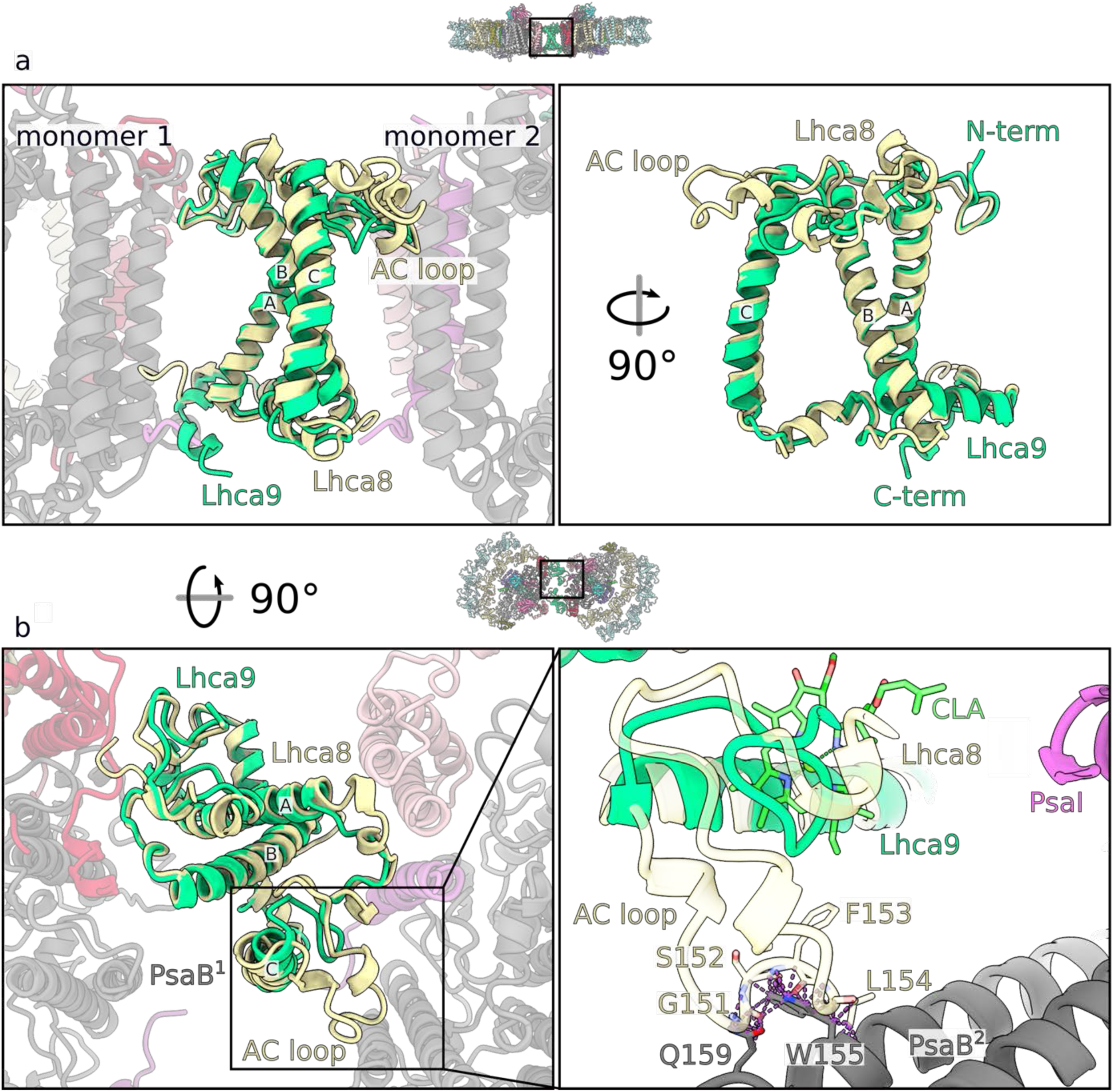
Lhca9 short AC-loop contributes to the dimerization. **a**, Superposition of Lhca8 and Lhca9, highlighting the difference in the AC-loop. **b**, The 90°-rotated view illustrates a clash between the longer AC-loop version and PsaB of monomer 2. Clashes between atoms with a VdW overlap >0.6 Å are highlighted by the dashed lines.

**Extended Data Fig. 6:**
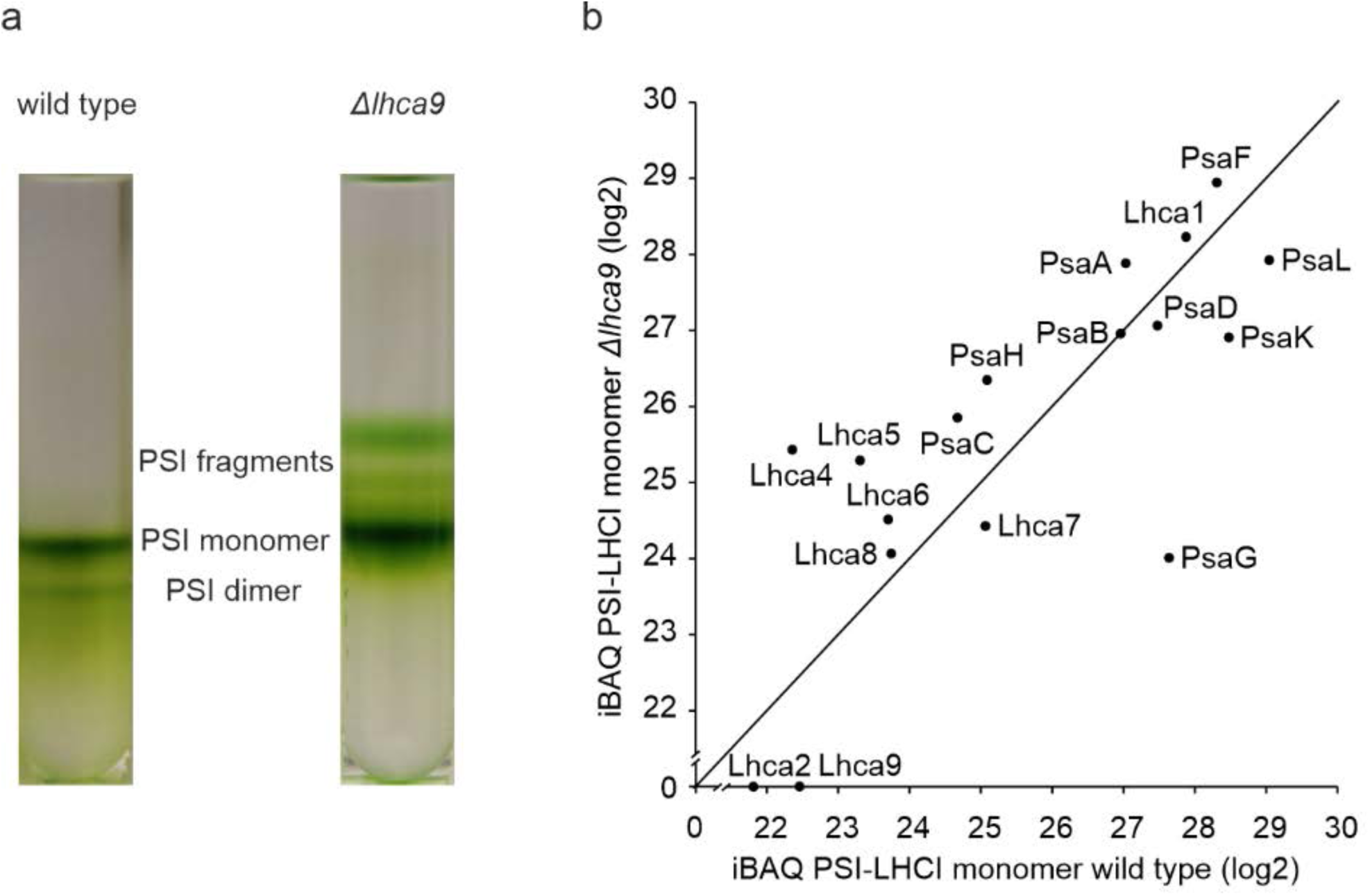
PSI dimer fraction is absent in *Δlhca9*. **a,** Sucrose density gradient of affinity purified PSI from *lhca9* insertional mutant (∼ 60 µg chl) compared to wild type. **b,** Quantitative mass spectrometry analysis of the PSI-LHC monomer fraction. iBAQ values are normalized to the PSI core subunit PsaB and values below 21 are excluded as they represent already low intensity values, which might not be reliable. PSI-LHCI subunits PsaE and Lhca3 were not detected at quantifiable intensities in this experiment. Detection of these small PSI and Lhca subunits via mass spectrometry depends on the presence of only a few proteotypic tryptic peptides, which could be missed, as measurements were done in a data dependent fashion (12 MS^2^ per MS^1^).

**Extended Data Fig. 7:**
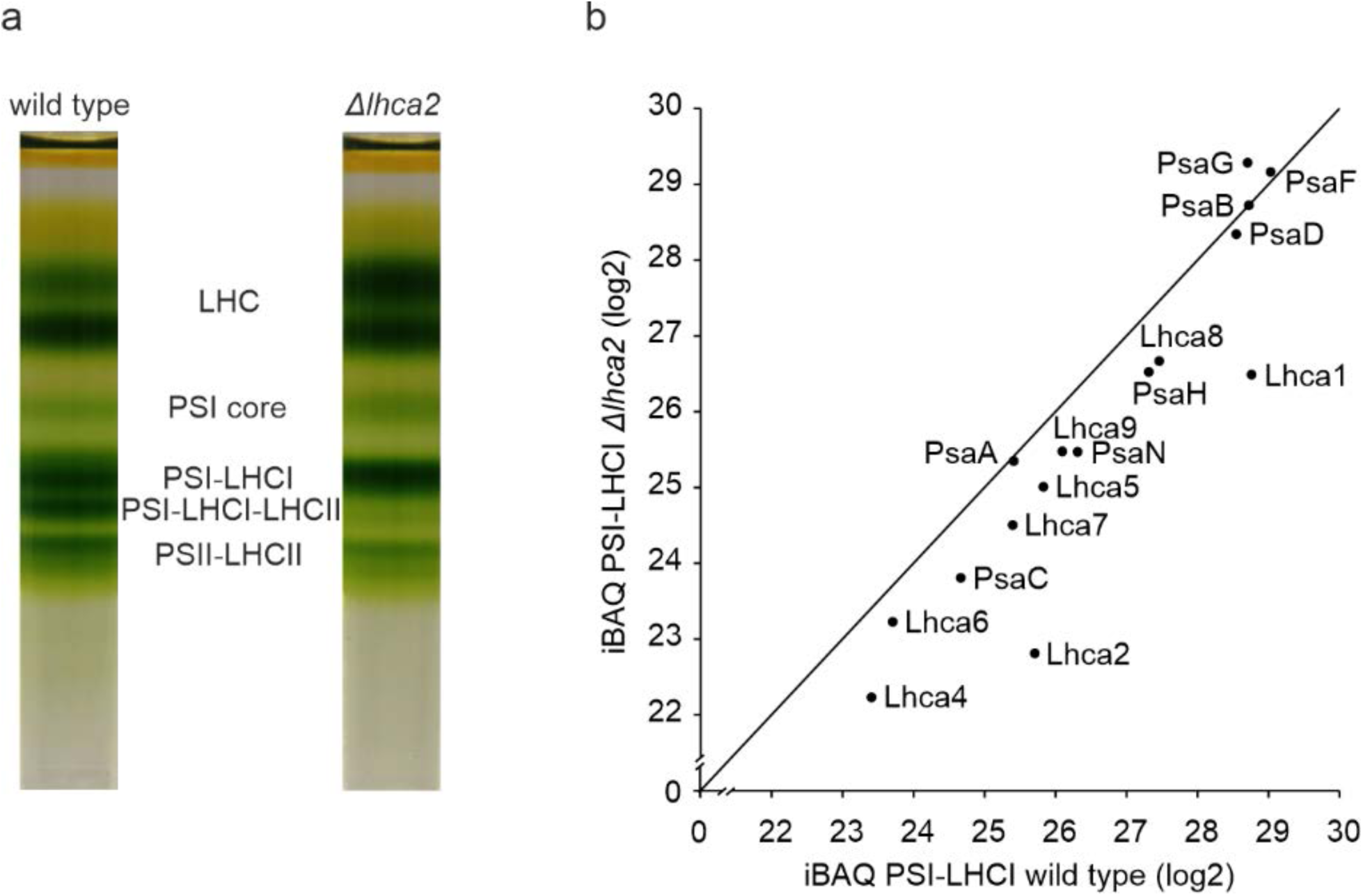
Lhca9 stably associates with PSI-LHCI despite reduction of Lhca2. **a,** Sucrose density gradients of α-DDM solubilized thylakoids (∼ 250 µg chl) isolated from wild type and *lhca2* insertional mutant. **b,** Quantitative mass spectrometry analysis of the PSI-LHCI fraction. iBAQ values are normalized to the PSI core subunit PsaB and values below 21 are excluded as they represent already low intensity values, which might not be reliable. PSI-LHCI subunits PsaE, PsaK and Lhca3 were not detected at quantifiable intensities in this experiment. Detection of these small PSI and Lhca subunits via mass spectrometry depends on the presence of only a few proteotypic tryptic peptides, which could be missed, as measurements were done in a data dependent fashion (12 MS^2^ per MS^1^).

**Extended Data Fig. 8:**
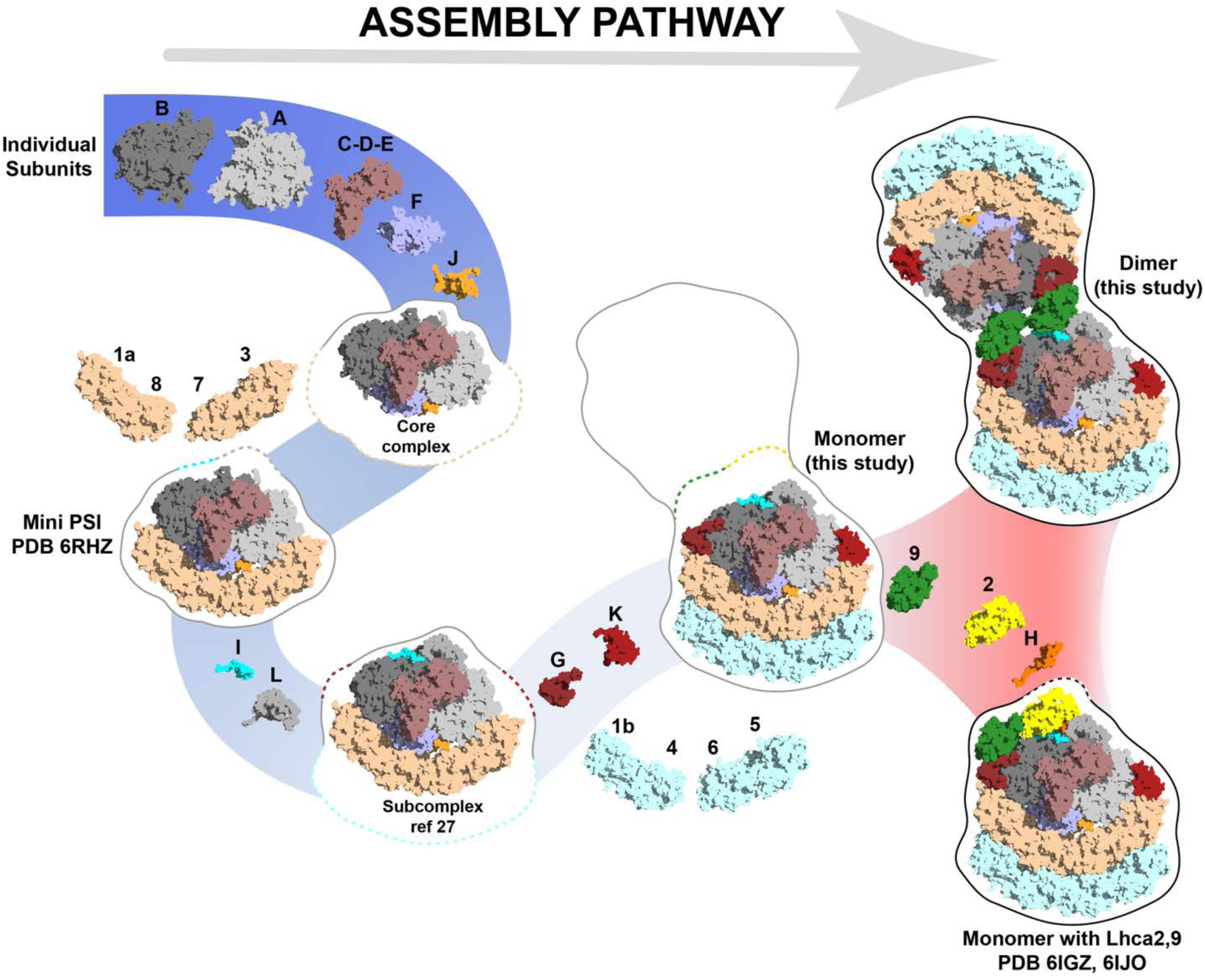
Assembly pathway of PSI towards dimer. **a**, Schematic view of known biochemically and structurally defined PSI states and assembling protein subunits. Subunit composition for each state is outlined with the shape of a following state and corresponding color. The divaricating of the assembly pathway takes place at the last stage (red background). The path towards PSI monomer or dimer is dependent on presence/absence of PsaH and Lhca2.

**Extended Data Fig. 9:**
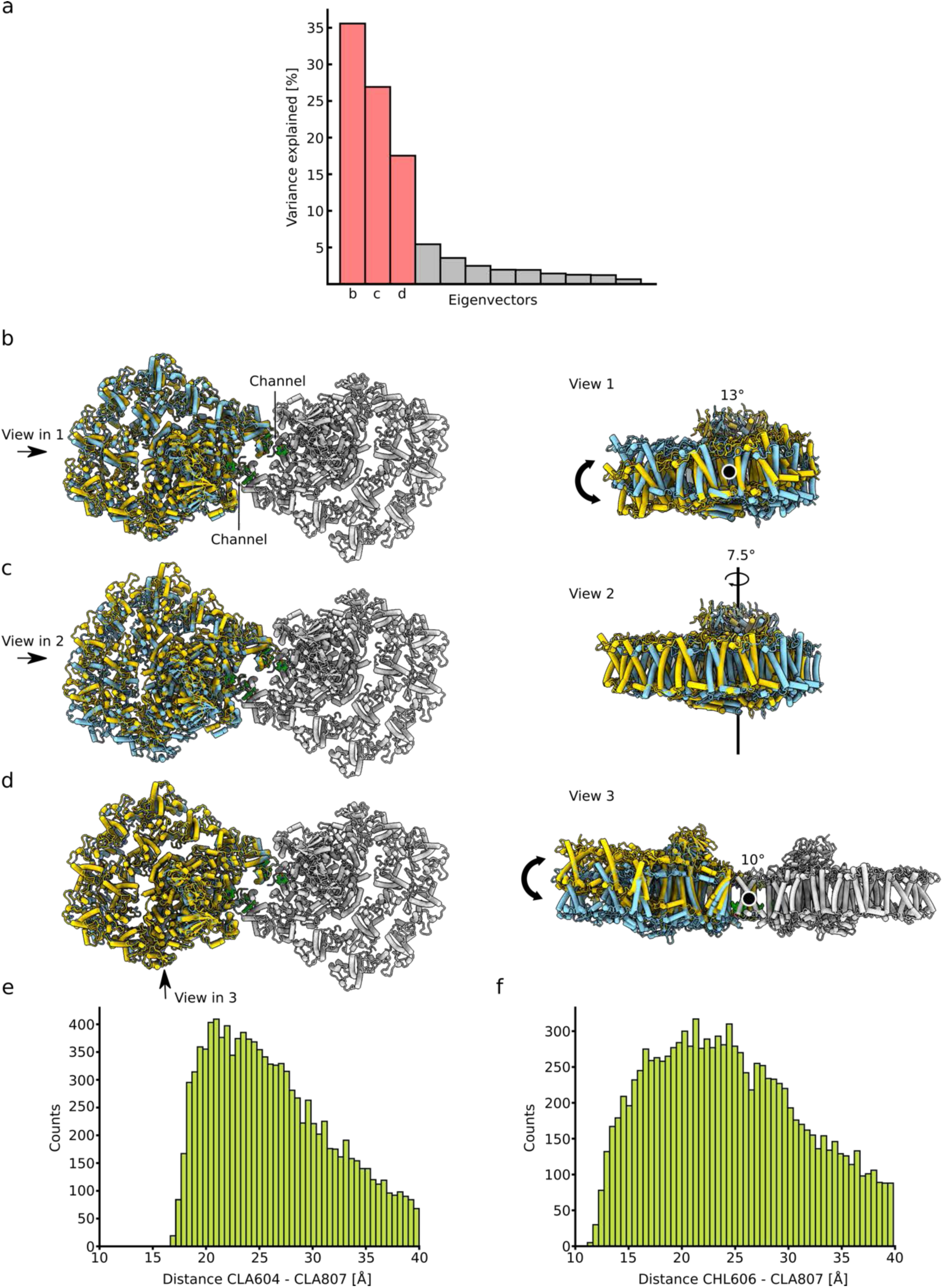
Multi-body refinement analysis. **a**, Eigenvectors that explain the variability of the data. The three major eigenvectors account for ∼78% of the motion in the PSI dimer. **b-d**, Stromal and side view of the model, showing the maximal motion along the three vectors. **e, f**, The distance between the chlorophylls is plotted based on the relative motion from the multi-body analysis. The y-axis shows the counts of particles that exhibit a certain distance.

**Extended Data Fig. 10:**
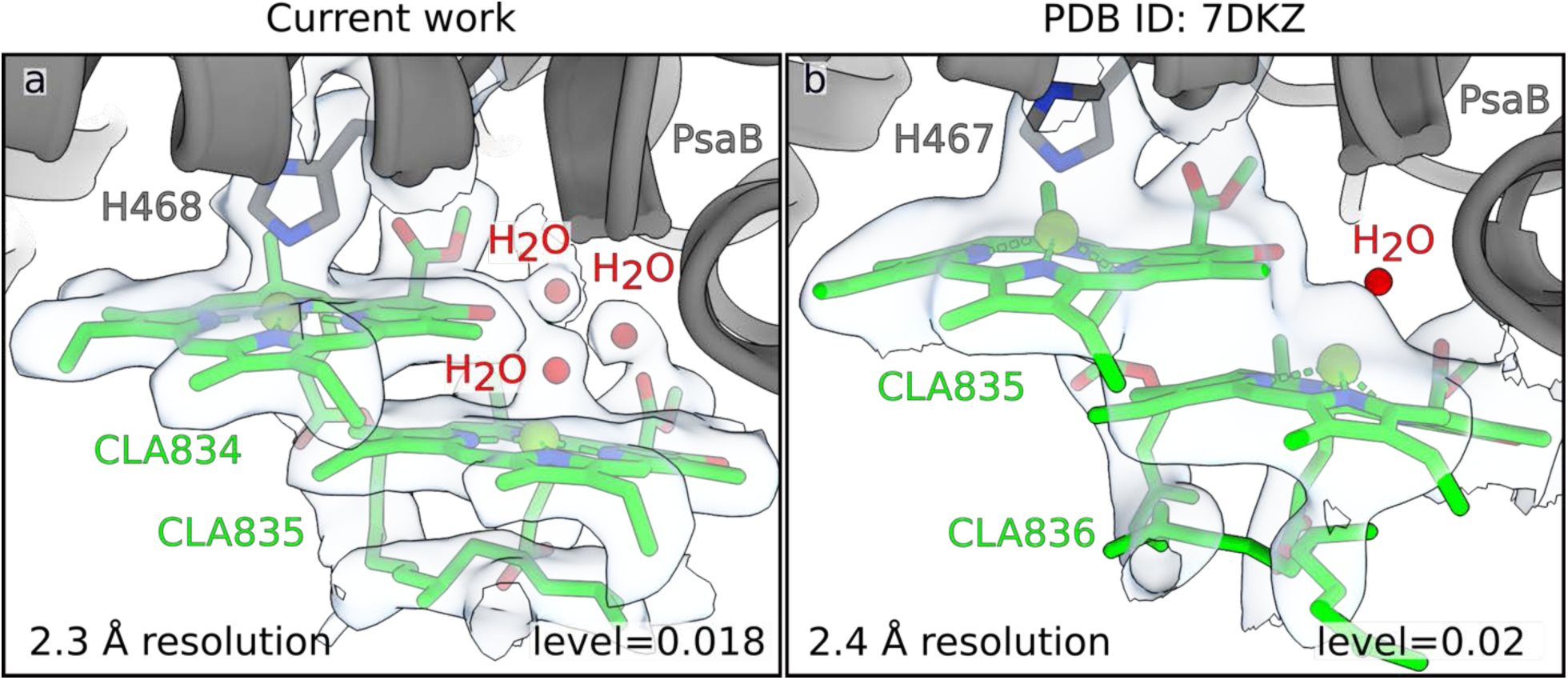
Comparison of the map/model quality with X-ray study. **a**, The current density around CLA834-835 is shown in iso-surface representation at a local resolution of 2.3 Å. Three water molecules involved in chlorophyll coordination are identified. Updated libraries are used for correct chlorophyll modeling. **b**, The same region from X-ray study at 2.4 Å resolution ^17^ lacks the density for waters, whereas the modeled water molecule has no density, and Mg atoms are not in the correct plane.

**Extended Data Fig. 11:**
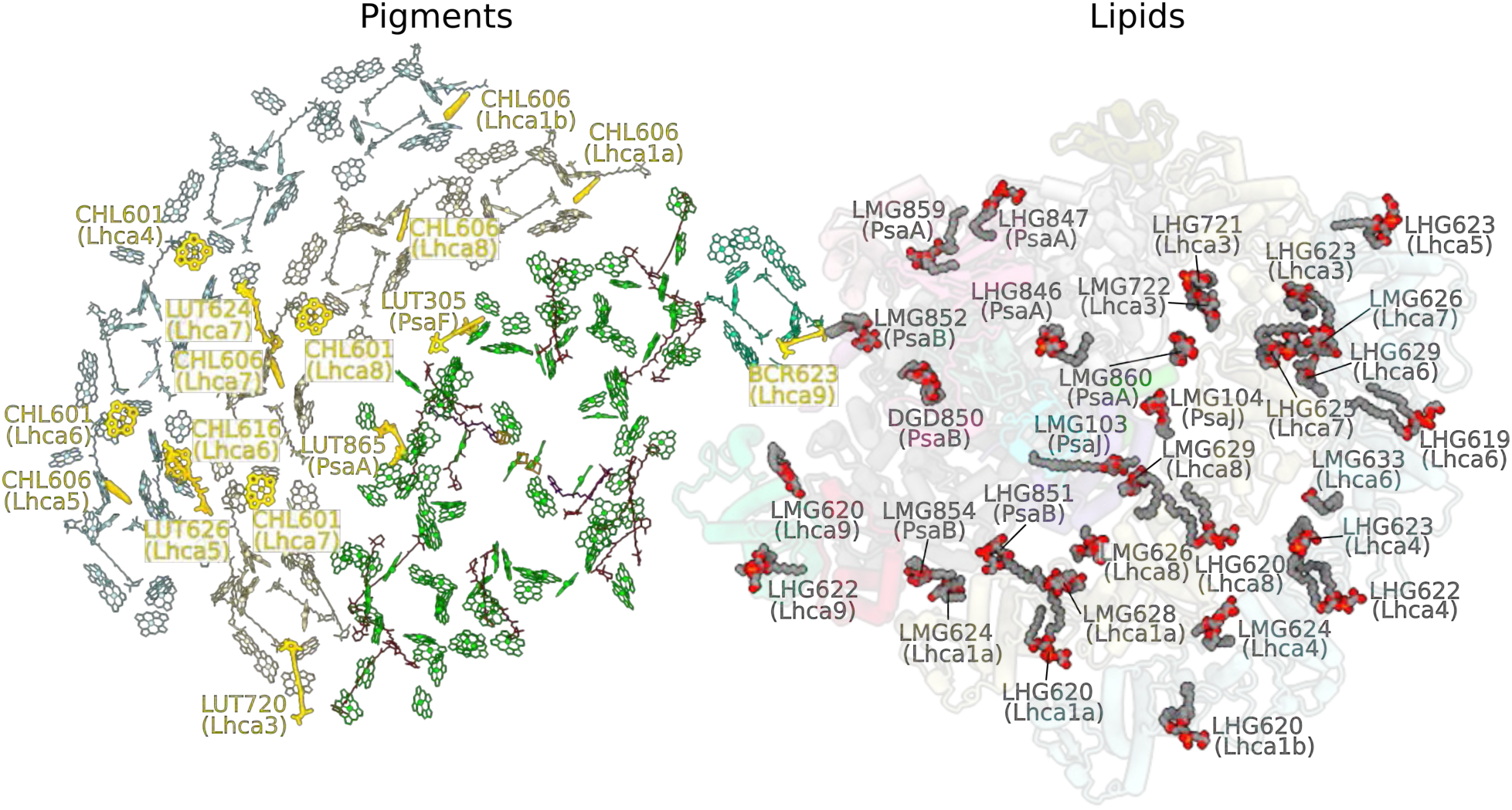
Overview of the pigments and lipids. Pigments (left) and lipids (right) are shown from stroma. The newly identified pigments are highlighted in bold gold.

**Extended Data Fig. 12:**
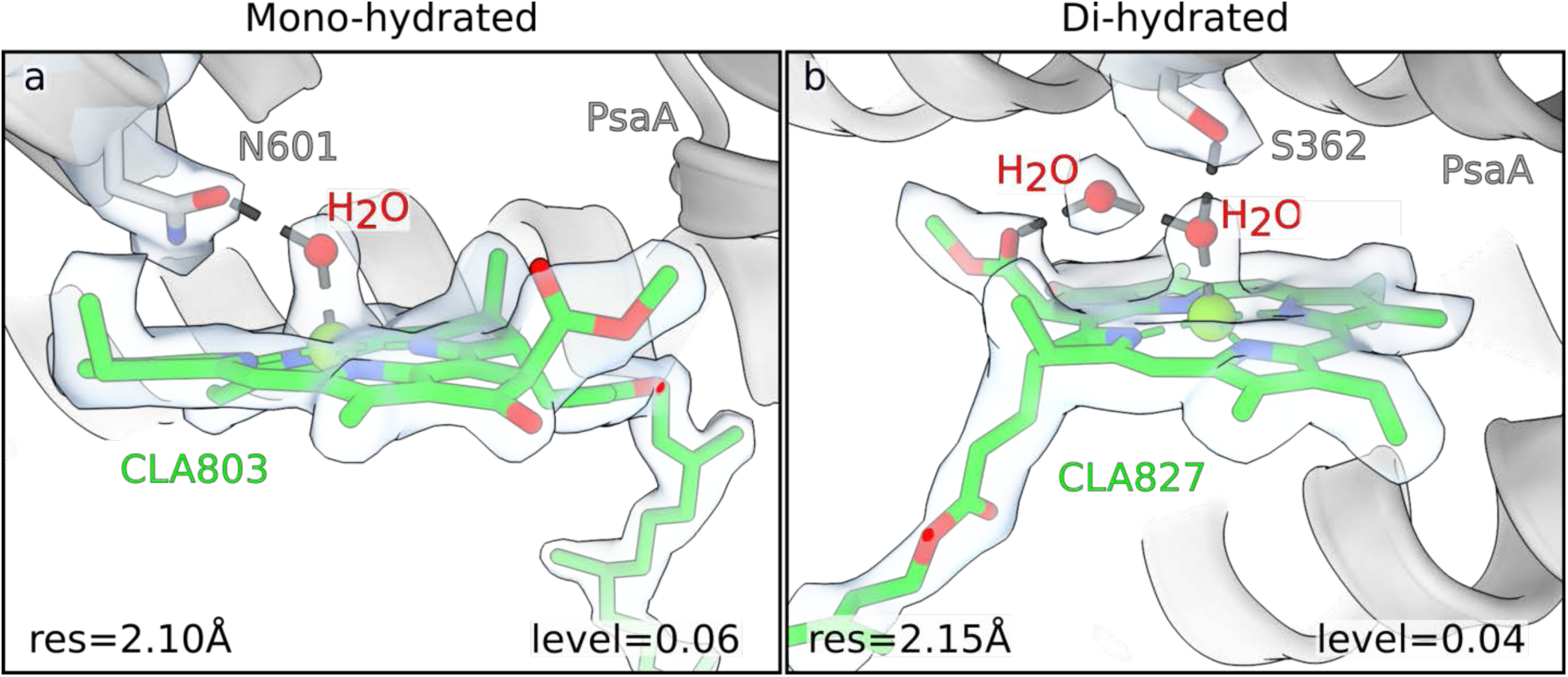
Mono- and di-hydrated chlorophylls. **a**, An example of a mono- hydrated chlorophyll with the corresponding density. **b,** An example of a di-hydrated chlorophyll with the corresponding density. Local resolution and map levels are indicated.

**Extended Data Table 1:** Protein composition of silver-stained spots from a 2D-PAGE of solubilized *C. reinhardtii* wild type thylakoids isolated from control and anoxic conditions.

| Spot | Identified proteins |
| --- | --- |
| control |  |
| 1 | Asa3, Asa4, Atp3 |
| 2 | $\beta$ mannosidase, Lhca3, Lhca5, Lhca6, Lhca7, LhcbM1 |
| 3 | $\beta$ mannosidase, PsbB, PsbC |
| 4 | Lhcb5, LhcbM3, Nad5, PsbB, PsbC |
| 5 | Lhcb4, Lhcb5, LhcbM4, PsbA, PsbB, PsbD |
| 6 | PsaA, PsaB |
| 7 | Asa4, AtpB, PsbA, PsbB, PsbC |
| 8 | LhcbM3, PsbA, PsbB, PsbC |
| 9 | Asa2, Lhca3, Lhca4, Lhca6, Lhcb5, LhcbM2, PsbB, PsbC |
| 10 | Aaa1, AtpB, LhcbM3, PsbA, PsbB, PsbC |
| 11 | LhcbM3, PsbB, PsbC |
| 12 | Lhcb4, Lhcb5, PetA, PsbA, PsbB, PsbC, PsbD |
| anoxia |  |
| 1 | Atp2, PsaA, PsaB |
| 2 | Lhcb5, LhcbM3, PsbA, PsbD |
| 3 | LhcbM3, LhcbM4, PsaB, PsbO |
| 4 | Lhca1, Lhca3, Lhca4, Lhca5, Lhca6, Lhca7, Lhca8 |
| 5 | Lhca3, Lhca5, Lhca6, Lhca7, Lhca8, Lhcb5, LhcbM1 |
| 6 | Lhca1, Lhca7, Lhca8, Lhca9, |
| 7 | Asa7, Lhca1, PsaB, PsaD |
| 8 | PsaF |
| 9 | PsaH |
| 10 | Asa2, PsbC |
| 11 | Asa6, PsaH |
| 12 | Lhcb4, Lhcb5, PsbA, PsbD |
| 13 | Nuo13, PsaF |
| 14 | AtpG |
| 15 | PetC |

**Supplementary Table 2:** Cryo-EM data collection, refinement and validation statistics of PSI monomer and dimer of *C. reinhartdii*.

|  | Dimer<br>(EMDB-xxxx)<br>(PDB xxxx) | Symmetry expanded<br>dimer<br>(EMDB-xxxx)<br>(PDB xxxx) | Monomer<br>(EMDB-xxxx)<br>(PDB xxxx) | Plastocyanin<br>(EMDB-xxxx)<br>(PDB xxxx) |
| --- | --- | --- | --- | --- |
| **Data collection and processing** |  |  |  |  |
| Magnification | 105,000 | 105,000 | 105,000 | 165,000 / 105,00 |
| Voltage (kV) | 300 | 300 | 300 | 300 |
| Electron exposure (e-/Å<sup>2</sup>) | 45.8 | 45.8 | 45.8 | 48 / 45.8 |
| Defocus range (µm) | -0.7-2.5 | -0.7-2.5 | -0.7-2.5 | -0.3, -1.1 / -0.7-2.5 |
| Pixel size (Å) | 0.84 | 0.84 | 0.84 | 0.51 / 0.84 |
| Initial particle images (no.) | 17,439 | 17,439 | 17,439 | 19,969 / 17,439 |
| Symmetry imposed | C2 | C1 | C1 | C1 |
| Final particle images (no.) | 14,173 | 28,346 | 74,209 | 66,080 |
| Map resolution (Å) | 2.97 | 2.74 | 2.31 | 2.55 |
| FSC threshold 0.143 |  |  |  |  |
| **Refinement** |  |  |  |  |
| Initial model used (PDB code) | 6JO5 | 6JO5 | 6JO5 | SWISS model |
| Map sharpening *B* factor (Å<sup>2</sup>) | -28.80 | -21.00 | -23.05 | -21.05 |
| Model composition |  |  |  |  |
| Non-hydrogen atoms | 100,830 | 56,460 | 50,499 | 729 |
| Protein residues | 8,418 | 4,712 | 4,164 | 98 |
| Ligands | 708 | 398 | 352 | 0 |
| Waters | 160 | 88 | 621 | 0 |
| *B* factors (Å<sup>2</sup>)<br>(min/max/mean) |  |  |  |  |
| Protein | 22.5/125.1/65.5 | 48.7/154.0/80.3 | 13.9/117.2/40.0 | 66.7/83.8/73.5 |
| Ligand | 25.5/118.9/66.9 | 51.0/148.5/79.45 | 15.9/109.5/39.45 |  |
| Waters | 25.4/115.3/64.0 | 51.1/95.5/72.0 | 13.4/57.5/31.72 |  |
| R.m.s. deviations |  |  |  |  |
| Bond lengths (Å) | 0.009 | 0.009 | 0.009 | 0.002 |
| Bond angles (°) | 1.065 | 1.069 | 1.085 | 0.494 |
| Validation |  |  |  |  |
| MolProbity score | 1.16 | 1.16 | 1.12 | 1.20 |
| Clashscore | 3.70 | 3.69 | 3.29 | 4.21 |
| Poor rotamers (%) | 0.39 | 0.35 | 0.24 | 0.00 |
| Ramachandran plot |  |  |  |  |
| Favored (%) | 98.40 | 98.34 | 97.98 | 98.96 |
| Allowed (%) | 1.57 | 1.61 | 1.99 | 1.04 |
| Disallowed (%) | 0.02 | 0.04 | 0.02 | 0.00 |

**Supplementary Table 3:** Chemical moieties involved in coordination of chlorophyll *a*/*b*.

|  | Chlorophyll *a/b* |  |  |  |  |  |  |  |  |  |  |
| --- | --- | --- | --- | --- | --- | --- | --- | --- | --- | --- | --- |
|  | His | Met | 1x H<sub>2</sub>O | 2x H<sub>2</sub>O | Gln | back-bone | Lipid | Asp | Glu | Asn | Total |
| **All subunits** | 84 | 2 | 13/1 | 22/19 | 13 | 11/7 | 10 | 5/3 | 28 | 11 | 199/30 |
| **PsaA** | 32 | 1 | 3 | 4 | 3 | 1 | 1 |  |  |  | 45 |
| **PsaB** | 31 | 1 | 5 |  | 1 |  | 1 | 1 |  |  | 40 |
| **PsaF** |  |  |  | 2 |  |  |  | 1 |  |  | 3 |
| **PsaG** | 1 |  |  |  |  |  |  | 1 |  |  | 2 |
| **PsaJ** |  |  |  |  |  |  |  |  | 1 |  | 1 |
| **PsaK** | 1 |  |  | 1 |  | 1 |  | 1 |  |  | 4 |
| **PsaL** | 1 |  | 1 |  |  |  |  |  |  |  | 2 |
| **Lhca1_a** | 2 |  |  | 3/2 |  | 1/1 | 1 |  | 3 | 1 | 11/3 |
| **Lhca1_b** | 2 |  | 1/1 | 2/1 |  | 1/1 | 1 |  | 3 | 1 | 11/3 |
| **Lhca3** | 2 |  | 1 | 2/1 | 2 | 2 |  |  | 3 | 1 | 13/1 |
| **Lhca4** | 1 |  |  | 1/3 | 1 | 0/1 | 1 | 1/1 | 3 | 2 | 10/5 |
| **Lhca5** | 3 |  |  | 2/3 | 1 | 2 | 1 | 0/1 | 3 | 1 | 13/4 |
| **Lhca6** | 3 |  | 1 | 1/3 | 1 | 0/2 | 1 | 0/1 | 3 | 1 | 11/6 |
| **Lhca7** | 2 |  | 1 | 2/2 | 1 | 1/1 | 1 |  | 3 | 1 | 12/3 |
| **Lhca8** | 2 |  |  | 2/2 | 1 | 1/1 | 1 |  | 3 | 1 | 11/3 |
| **Lhca9** | 1 |  |  | 0/2 | 2 | 1 | 1 |  | 3 | 2 | 10/2 |

